# NLRC5 regulates expression of MHC-I and provides a target for anti-tumor immunity in transmissible cancers

**DOI:** 10.1101/2020.09.06.274720

**Authors:** Chrissie E. B. Ong, Amanda L. Patchett, Jocelyn M. Darby, Jinying Chen, Guei-Sheung Liu, A. Bruce Lyons, Gregory M. Woods, Andrew S. Flies

## Abstract

Downregulation of major histocompatibility complex I (MHC-I) on tumor cells is a primary means of immune evasion by many types of cancer. Additionally, MHC-I proteins are a primary target of immune-mediated transplant rejection. Transmissible tumors that overcome allograft rejection mechanisms and evade anti-tumor immunity have killed thousands of wild Tasmanian devils (*Sarcophilus harrisii*). Interferon gamma (IFNG) upregulates surface MHC-I expression on devil facial tumor (DFT) cells but is not sufficient to induce tumor regressions. Transcriptome analysis of IFNG-treated DFT cells revealed strong upregulation of *NLRC5*, a master regulator of MHC-I in humans and mice. To explore the role of NLRC5 in transmissible cancers, we developed DFT cell lines that constitutively overexpress NLRC5. Transcriptomic results suggest that the role of NLRC5 as a master regulator of MHC-I is conserved in devils. Furthermore, NLRC5 was shown to drive the expression of many components of the antigen presentation pathway. To determine if MHC-I is a target of allogeneic immune responses, we tested serum from devils with anti-DFT responses including natural DFT regressions against DFT cells. Antibody binding occurred with cells treated with IFNG and overexpressed NLRC5. However, CRISPR/Cas9-mediated knockout of MHC-I subunit beta-2-microglobulin (*B2M*) eliminated antibody binding to DFT cells. Consequently, MHC-I could be identified as a target for anti-tumor and allogeneic immunity and provides mechanistic insight into MHC-I expression and antigen presentation in marsupials. NLRC5 could be a promising target for immunotherapy and vaccines to protect devils from transmissible cancers and inform development of transplant and cancer therapies for humans.

## INTRODUCTION

In 1996, a wild Tasmanian devil (*Sarcophilus harrisii*) was photographed with a large facial tumor. In subsequent years, similar devil facial tumors (DFTs) were recorded^1^, and in 2006, it was confirmed that DFTs are clonally transmissible cancers that spread among devils through social interactions^2,3^. In 2014, a second genetically independent transmissible devil facial tumor (DFT2) was discovered in southern Tasmania^4^. Despite the independent origin of the first devil facial tumor (DFT1) and DFT2, both clonal tumors arose from a Schwann cell lineage^5,6^, suggesting devils could be prone to transmissible Schwann cell cancers. These lethal and unique tumors are simultaneously cancers, allografts, and infectious diseases, and have been the primary driver of an average 77% decline in devil populations across the island state of Tasmania^7^.

The successful transmission and seeding of DFT cells from one devil to another as an allograft^3^ reveals its ability to circumvent both allogeneic and anti-tumor immune responses. DFT1 cells generally express little or no major histocompatibility complex class I (MHC-I) on their surface^8^, an immune escape mechanism commonly observed in human cancers^9^ that prevents recognition of tumor cells by cytotoxic anti-tumor CD8^+^ T cells. Beta-2-microglobulin (B2M) is necessary for surface MHC-I expression and the clonal DFT1 cell lineage has a hemizygous mutation in the *B2M* gene^10^, suggesting that immune evasion through reduced MHC-I expression has been a target of evolutionary selection pressure. Loss of MHC-I should lead to recognition and cytotoxic responses by natural killer (NK) cells. Devils have demonstrated NK-like activity *in vitro*^11^ but the ongoing transmission of DFT1 cells suggests that NK cytotoxic response against DFT1 cells either do not occur or are ineffective. All DFT1 cell lines tested to date can upregulate MHC-I in response to interferon gamma (IFNG) treatment^8^. Rare cases of DFT1 regression have been reported in the wild^12^ and serum antibody responses of these devils are generally higher against cell lines treated with IFNG to upregulate MHC-I^12,13^. In contrast to DFT1 cells, DFT2 cells constitutively express MHC-I, but the most highly-expressed alleles appear to be those shared by the DFT2 cells and the host devil^14^. This further suggest a critical role of MHC-I in immune evasion by DFT cells.

Upregulation of MHC-I on DFT1 cells via treatment with IFNG has served as the foundation for a vaccine against devil facial tumor disease (DFTD), which is caused by DFT1 cells. However, there are caveats to using a pleiotropic cytokine such as IFNG. IFNG plays multiple roles in the innate and adaptive immune system and can function to drive either an anti-tumor or a pro-tumor response depending on the circumstances^15^. While IFNG is well known for directing the immune response towards anti-tumor immunity, it also causes the upregulation of programmed death ligand 1 (PDL1)^16^ and non-classical, monomorphic MHC-I SAHA-UK on DFT cells^14^. PDL1 and SAHA-UK molecules can be counterproductive to the cell-mediated immune response mediated by MHC-I recognition. Additionally, the inhibition of cell proliferation and increased DFT cell death associated with IFNG^17^ constrain large-scale production of IFNG-treated DFT cells for whole cell vaccines.

NLRC5 (NLR caspase recruitment domain containing protein 5), a member of the NOD-like receptor (NLR) family, was identified in 2010 as the transcriptional activator of MHC-I genes^18^. NLRC5 is strongly upregulated by IFNG and is found to be a critical mediator for IFNG-induced MHC-I expression in humans and mice^18^, but little is known about NLRC5 in other species. NLRC5 acts with high specificity^18^, and functions in MHC-I regulation by interacting with several other transcription factors^19^ to form a multi-protein complex called the enhanceosome^20,21^. The enhanceosome activates the promoters of MHC-I genes and components of the antigen processing machinery such as B2M, immunoproteasome subunits PSMB8 (also known as LMP7) and PSMB9 (also known as LMP2), and transporter associated with antigen processing 1 (TAP1)^9,18^. Aside from MHC-I regulation, NLRC5 has been reported to be involved in innate immune responses as well as malignancy of certain cancers^22^. Despite a potential central role of NLRC5 in immune evasion, studies of NLRC5 remain limited and several hypothesized secondary roles of NLRC5 remain unexplored^22^.

In this study, we take advantage of a unique natural experiment in which two independent clonal tumor cell lines have essentially been passaged through hundreds of free-living animals to assess the role of NLRC5 and MHC-I in immune evasion. The overexpression of NLRC5 in DFT1 and DFT2 cells induced the expression of *B2M,* MHC-I heavy chain *SAHAI-01* and other functionally-related genes. *PDL1* and the non-classical MHC-I *SAHA-UK* which are upregulated by IFNG were not induced by NLRC5. MHC-I was constitutively expressed on the surface of DFT cells overexpressing NLRC5, which suggests that modulation of NLRC5 expression could be a potential substitute for IFNG to increase DFT cell immunogenicity. Additionally, MHC-I molecules on DFT cells were revealed to be an immunogenic target of allogeneic responses in wild devils.

## MATERIALS AND METHODS

### Cells and Cell Culture Conditions

DFT1 cell line C5065 strain 3 ^23^ (RRID:CVCL_LB79) and DFT2 cell lines RV (RRID:CVCL_LB80) and JV (RRID not available) were used in this study as indicated. DFT1 C5065 was provided by A-M Pearse and K. Swift of the Department of Primary Industries, Parks, Water and Environment (DPIPWE) (Hobart, TAS, Australia) and was previously established from DFT1 biopsies obtained under the approval of the Animal Ethics Committee of the Tasmanian Parks and Wildlife Service (permit numbers 33/2004-5 and 32/2005-6). DFT2 cell lines RV and JV were established from single cell suspensions obtained from tumor biopsies^4^. Cells were cultured at 35 °C with 5% CO_2_ in complete RPMI medium: RPMI 1640 medium with L-glutamine (Thermo Fisher Scientific, Waltham, MA, USA), 10% heat-inactivated fetal bovine serum (Bovogen Biologicals, Melbourne, VIC, Australia), 1% (v/v) Antibiotic-Antimycotic (100X) (Thermo Fisher Scientific), 10 mM HEPES (Thermo Fisher Scientific) and 50 μM 2-mercaptoethanol (Sigma-Aldrich, St. Louis, MO, USA).

### RNA Sequencing and Analysis

Initial RNA sequencing was performed using DFT1 C5065 and DFT2 RV cells treated with and without 5 ng/mL recombinant devil IFNG (provided by Walter and Eliza Hall Institute (WEHI), Melbourne, VIC, Australia) for 24 h according to the previously described protocols^6,24^. For the remaining cell lines (**Table 1**, ID # 5–9), total RNA was extracted using the NucleoSpin® RNA plus kit (Macherey Nagel, Düren, Germany) per manufacturer’s instructions. Two replicates were prepared for each cell line. RNA sequencing was conducted at the Ramaciotti Centre for Genomics (Sydney, NSW, Australia) using the following methods. RNA integrity was assessed using Agilent TapeStation (Agilent Technologies, Santa Clara, CA, USA). All samples had RNA Integrity Number (RIN) scores of 10.0. mRNA libraries were prepared using the TruSeq Stranded mRNA Library Prep (Illumina Inc., San Diego, CA, USA). The libraries were sequenced on an Illumina NovaSeq 6000 platform (Illumina) with 100 base-pair single-end reads. The quality of the sequencing reads were analyzed using FastQC version 0.11.9 ^25^. Raw FASTQ files have been deposited to the European Nucleotide Archive (ENA) and are available at BioProject # PRJEB39847.

**Table 1.** Devil facial tumor (DFT) cell lines and treatments

| ID # | Sample name | Parent cell line | Treatment |
| --- | --- | --- | --- |
| 1 | DFT1.WT* | DFT1 C5065 | Untreated |
| 2 | DFT1.WT + IFNG | DFT1 C5065 | 5 ng/mL IFNG, 24h |
| 3 | DFT2.WT<sup>RV</sup>* | DFT2 RV | Untreated |
| 4 | DFT2.WT<sup>RV</sup> + IFNG | DFT2 RV | 5 ng/mL IFNG, 24h |
| 5 | DFT1.BFP | DFT1 C5065 | Transfected with control vector pSBbi-BH |
| 6 | DFT1.NLRC5 | DFT1 C5065 | Transfected with NLRC5 vector pCO1 |
| 7 | DFT2.WT | DFT2 JV | Untreated |
| 8 | DFT2.BFP | DFT2 JV | Transfected with control vector pSBbi-BH |
| 9 | DFT2.NLRC5 | DFT2 JV | Transfected with NLRC5 vector pCO1 |
| 10 | DFT1.B2M<sup>-/-</sup> | DFT1 C5065 | Transfected with *B2M* targeting vector pAF217 |
| 11 | DFT1.B2M<sup>-/-</sup> + IFNG | DFT1 C5065 | Transfected with *B2M* targeting vector pAF217 and treated with 5 ng/mL IFNG for 24h |
| 12 | DFT1.NLRC5.B2M<sup>-/-</sup> | DFT1 C5065 | Transfected with NLRC5 vector pCO1 and *B2M* targeting vector pAF218 |

\*DFT1.WT data from Patchett et al., 2018<sup>24</sup> and DFT2.WT<sup>RV</sup> from Patchett et al., 2020<sup>6</sup> available through European Nucleotide Archive # PRJNA416378 and # PRJEB28680, respectively.

The sequencing reads were mapped to the Tasmanian devil reference genome (GCA_902635505.1 mSarHar1.11) using Subread version 2.0.0 ^26^. Uniquely mapped reads were counted and assigned to genes using featureCounts^27^. Differential expression analysis of gene counts was performed using statistical software R studio^28^ on R version 4.0.0 ^29^. Firstly, genes with less than 100 aligned reads across all samples were filtered out to exclude lowly expressed genes. Gene counts were then normalized across samples by upper quartile normalization using edgeR^30-32^ and EDASeq^33,34^. Normalized read counts were scaled by transcripts per kilobase million (TPM) to account for varied gene lengths. For differential expression analysis, gene expression of NLRC5-overexpressing cell lines (DFT1.NLRC5, DFT2.NLRC5) were compared against BFP-control cell lines (DFT1.BFP, DFT2.BFP) while IFNG-treated cells (DFT1.WT + IFNG, DFT2.WT^RV^ + IFNG) were compared against the untreated wild-type (DFT1.WT, DFT2.WT^RV^), according to their respective tumor origin. Differential gene expression was calculated using the *voom*^35^ function in *limma*^36^ with linear modelling and empirical Bayes moderation^37^ (**Supplementary Table 1**). Genes were defined as significantly differentially expressed by applying FDR < 0.05, and log_2_ fold change (FC) ≥ 2.0 (upregulated) or ≤ −2.0 (downregulated) thresholds.

A bar plot of fold change in mRNA expression upon treatment was created from TPM values in GraphPad Prism version 5.03. Venn diagrams of differentially expressed genes were developed using Venny version 2.1 ^38^. Heatmaps were created from log_2_TPM values using the ComplexHeatmap^39^ package in R studio. For functional enrichment analysis, over-representation of gene ontology (GO) and Reactome pathways was analyzed on differentially expressed genes in R studio using functions *enrichGO* in ClusterProfiler^40^ and *enrichPathway* in ReactomePA^41^, respectively. Significant GO terms and Reactome pathways were selected by applying the cut-offs p-value < 0.001, q-value < 0.05 and adjusted p-value < 0.05. P-values were adjusted for multiple testing using Benjamini-Hochberg method.

### Plasmid Construction

The coding sequence for full length devil *NLRC5* (ENSSHAT00000015489.1) was isolated from cDNA of devil lymph node mononuclear cells stimulated with recombinant devil IFNG^16^ (10 ng/mL, 24 h). Devil *NLRC5* was then cloned into plasmid pAF105 (detailed description of pAF105 plasmid construction available in **Supplementary Methods 1**). For this study, devil *NLRC5* was amplified from pAF105 with overlapping ends to the 5’ and 3’ SfiI sites of the Sleeping Beauty transposon plasmid pSBbi-BH^42^ (a gift from Eric Kowarz; Addgene # 60515, Cambridge, MA, USA) using Q5® Hotstart High-Fidelity 2X Master Mix (New England Biolabs (NEB), Ipswich, MA, USA) (see **Supplementary Table 2** for primers and reaction conditions). The fragment was cloned into SfiI-digested (NEB) pSBbi-BH using NEBuilder® HiFi DNA Assembly Cloning Kit (NEB) and the assembled plasmid pCO1 was transformed into NEB® 5-alpha competent *Escherichia coli* (High Efficiency) (NEB) according to manufacturer’s instructions (see **Supplementary Figure 1** for plasmid maps). Positive clones were identified by colony PCR and the plasmid was purified using NucleoSpin^®^ Plasmid EasyPure kit (Macherey-Nagel). The cloned devil *NLRC5* transcript was verified by Sanger sequencing using Big Dye™ Terminator v3.1 Cycle Sequencing Kit (Applied Biosystems (ABI), Foster City, CA, USA) and Agencourt® CleanSEQ® (Beckman Coulter, Brea, CA, USA) per manufacturer’s instructions. The sequences were analyzed on 3500xL Genetic Analyzer (ABI) (see **Supplementary Table 3** for list of sequencing primers). For detailed step-by-step protocols for plasmid design and construction, reagent recipes, and generation of stable cell lines, see Bio-protocol # e3986^43^.

### Transfection and Generation of Stable Cell Lines

Stable cell lines of both DFT1 and DFT2 (C5065 and JV cell lines respectively) overexpressing NLRC5 were prepared as follows. 5×10^5^ cells were seeded into a 6-well plate and incubated overnight to achieve 50-80% confluency on the day of transfection. As the vector constructed uses a Sleeping Beauty (SB) transposon system for gene transfer, co-transfection of an expression vector encoding a SB transposase enzyme pCMV(CAT)T7-SB100^44^ (a gift from Zsuzsanna Izsvak; Addgene plasmid # 34879) was needed to facilitate this process. Per 2.0 mL of culture volume, 2.0 μg of plasmid DNA (1.5 μg pCO1 + 0.5 μg pCMV(CAT)T7-SB100) was diluted in phosphate-buffered saline (PBS) to 100 μL and then added to 6.0 μg of polyethylenimine (PEI) (1 mg/mL, linear, 25 kDa; Polysciences, Warrington, FL, USA) diluted in PBS to 100 μL (3:1 ratio of PEI to DNA (w/w)). The DNA:PEI solution was mixed by gentle pipetting and incubated at room temperature for 15 to 20 min. The media on DFT cells were replaced with fresh complete RPMI medium and the transfection mix was added dropwise to the cells. The cells were incubated with the DNA:PEI solution overnight at 35 °C with 5% CO_2_. The next morning, media was replaced with fresh complete RPMI medium. 48 h post-transfection, the cells were observed for fluorescence through expression of reporter gene mTagBFP and were subjected to seven days of positive selection by adding 1 mg/mL hygromycin B (Sigma-Aldrich) in complete RPMI medium. Once selection was complete, the cells were maintained in 200 μg/mL hygromycin B in complete RPMI medium. pSBbi-BH was used as a control to account for the effects of the transfection and drug selection process.

### Flow Cytometric Analysis of B2M Expression

Cells were harvested and plated in a round-bottom 96-well plate (1×10^5^ per well) and centrifuged at 500*g* for 3 min at 4 °C to discard the medium. Cells were blocked with 50 μL of 1% normal goat serum (Thermo Fisher Scientific) in FACS buffer (PBS with 0.5% BSA, 0.02% sodium azide) for 10 min on ice. After blocking, 0.4 μL anti-devil B2M mouse antibody in supernatant (13-34-45, a gift from Hannah Siddle)^8^ diluted to a total of 50 μL in FACS buffer was added to the cells for 15 min on ice. The cells were washed with 150 μL FACS buffer and centrifuged at 500*g* for 3 min at 4 °C. Goat anti-mouse IgG-Alexa Fluor 488 (Thermo Fisher Scientific) was diluted in FACS buffer to 4 μg/mL and 50 μL of the solution was incubated with the target cells in the dark for 30 min on ice. The cells were washed twice with FACS buffer to remove excess secondary antibody. Lastly, the cells were resuspended in 200 μL FACS buffer with propidium iodide (PI) (500 ng/mL) (Sigma-Aldrich) prior to analysis on BD FACSCanto™ II (BD Biosciences, Franklin Lakes, NJ, USA). As a positive control for surface B2M expression, DFT1 C5065 and DFT2 JV cells were stimulated with 5 ng/mL recombinant devil IFNG^16^ for 24 h.

### Generation of *B2M* CRISPR/Cas9 Knockout Cell Lines (B2M^-/-^)

Two single guide RNAs (sgRNAs) targeting the first exon of devil *B2M* gene (ENSSHAG00000017005) were designed using a web-based CRISPR design tool CHOPCHOP^45^ (**Supplementary Figure 2**). Complementary oligonucleotides encoding each *B2M* sgRNA sequence were synthesized (Integrated DNA Technologies (IDT), Coralville, IA, USA), phosphorylated and annealed before cloning into lentiCRISPRv2 plasmid^46^ (a gift from Feng Zhang; Addgene # 52961) at BsmBI (NEB) restriction sites using T4 DNA ligase (NEB) (see **Supplementary Table 4** for oligonucleotide sequences). The ligated plasmids pAF217 and pAF218 were then transformed into NEB® Stable Competent *Escherichia coli* (High Efficiency) (NEB). Single colonies were selected, and the plasmids were purified using ZymoPURE™ Plasmid Miniprep Kit (Zymo Research, Irvine, CA, USA). The sgRNA sequence in each plasmid was validated by Sanger sequencing according to the method described above (see **Supplementary Table 3** for list of sequencing primers).

B2M targeting vectors pAF217 and pAF218 were each transfected into DFT1.WT and DFT1.NLRC5 cells to generate *B2M* knockout cell lines DFT1.B2M^-/-^ and

DFT1.NLRC5.B2M^-/-^. Transfection of cells were carried out as described above with the exception that 1.5 μg of plasmid was used instead of 2.0 μg. A day after transfection, the cells were subjected to positive selection by adding 100 μg/mL puromycin (InvivoGen, San Diego, CA, USA) for a week.

Post-drug selection, the cells were screened and sorted multiple rounds using a Beckman-Coulter MoFlo Astrios cell sorter to select DFT1.B2M^-/-^ and DFT1.NLRC5.B2M^-/-^ cells with negative B2M expression. DFT1.B2M^-/-^ cells were treated with 10 ng/mL devil recombinant IFNG^16^ for 24 h to stimulate surface B2M upregulation prior to analysis. For flow cytometry, cells were first harvested by centrifugation at 500*g* for 3 min at 4 °C, and then blocked with 100 μL of 1% normal goat serum (Thermo Fisher Scientific) in complete RPMI medium for 10 min on ice. After blocking, the cells were incubated with 0.8 μL anti-devil B2M mouse antibody in supernatant^8^ diluted in complete RPMI to a total of 100 μL for 15 min on ice. The cells were washed with 2.0 mL complete RPMI and centrifuged at 500*g* for 3 min at 4 °C. Next, the cells were incubated with 100 μL of 2 μg/mL goat anti-mouse IgG-Alexa Fluor 647 (Thermo Fisher Scientific) diluted in complete RPMI in the dark for 15 min on ice. The cells were washed with 2.0 mL of complete RPMI medium to remove excess secondary antibody. Lastly, the cells were resuspended to a concentration of 1×10^7^ cells/ml in 200 ng/mL DAPI (Sigma-Aldrich) diluted in complete RPMI medium. B2M negative cells were selected and bulk-sorted using cell sorter Moflo Astrios EQ (Beckman Coulter).

After multiple rounds of sorting to establish a B2M negative population, genomic DNA of the cells was isolated and screened for mutations in the *B2M* gene by Sanger sequencing (see **Supplementary Table 3** for sequencing primers). Indels (insertions or deletions) in the *B2M* gene were assessed using Inference of CRISPR Edits (ICE) analysis tool version 2.0 from Synthego^47^ (Menlo Park, CA, USA) (**Supplementary Figure 2**). *B2M* knockout cell lines: (i) DFT1.B2M^-/-^ derived from DFT1 cells transfected with pAF217, and (ii) DFT1.NLRC5.B2M^-/-^ derived from DFT1.NLRC5 transfected with pAF218 were selected for downstream analysis (see **Table 1** for full list of cell lines).

### Flow Cytometric Analysis of Serum Antibody Target

Serum samples from wild Tasmanian devils were collected as described^12,48^. To induce surface expression of MHC-I, DFT cells were treated with 10 ng/mL devil recombinant IFNG^16^ for 24 h prior to analysis. Cells were washed with cold FACS buffer and 1×10^5^ cells per well were plated in a round-bottom 96-well plate. The cells were centrifuged at 500*g* for 3 min at 4 °C to discard the medium. Serum samples (see **Supplementary Table 5** for serum sample information) were thawed on ice and diluted 1:50 with FACS buffer. 50 μL of diluted serum was added to the cells and incubated for 1 h on ice. After incubation, the cells were washed twice with 200 μL FACS buffer. 50 μL of 10 μg/mL monoclonal mouse anti-devil IgG2b antibody (A4-D1-2-1, provided by WEHI)^49^ in FACS buffer was added to the cells and incubated for 30 min on ice. The cells were washed twice with FACS buffer and then incubated with 50 μL of 4 μg/ml goat anti-mouse IgG-Alexa Fluor 488 (Thermo Fisher Scientific) in FACS buffer for 30 min on ice, protected from light. The cells were washed twice with ice-cold PBS (Thermo Fisher Scientific). After washing, the cells were stained with LIVE/DEAD™ Fixable Near-IR Dead Cell Stain (Thermo Fisher Scientific) per manufacturer’s instructions. For B2M surface expression analysis, the cells were stained as described in the protocol above. However, LIVE/DEAD™ Fixable Near-IR Dead Cell Stain (Thermo Fisher Scientific) was used instead of PI to determine cell viability. All cells were fixed with FACS fix (0.02% sodium azide, 1.0% glucose, 0.4% formaldehyde) diluted by 20 times prior to analysis on BD FACSCanto™ II (BD Biosciences).

## RESULTS

### NLRC5 is upregulated in DFT1 and DFT2 cells treated with IFNG

IFNG has been shown to upregulate MHC-I^8^ and PDL1^16^ on DFT cells. To probe the mechanisms driving upregulation of these key immune proteins, we performed RNA-seq using mRNA extracted from IFNG-treated DFT1 cell line C5065 (DFT1.WT) and an IFNG-treated DFT2 cell line RV (DFT2.WT^RV^). Markers for Schwann cell differentiation, SRY-box 10 (SOX10) and neuroepithelial marker nestin (NES), that are expressed in both DFT1 and DFT2 cells^6^, were selected as internal gene controls. As expected, transcriptome analysis showed that *B2M*, MHC-I gene *SAHAI-01*, and *PDL1* were strongly upregulated by IFNG. MHC-I transactivator *NLRC5* was also upregulated upon IFNG treatment, more than a 100-fold in DFT1.WT (275-fold) and DFT2.WT^RV^ cells (124-fold) relative to untreated cells (**Fig. 1**).

**Figure 1.**
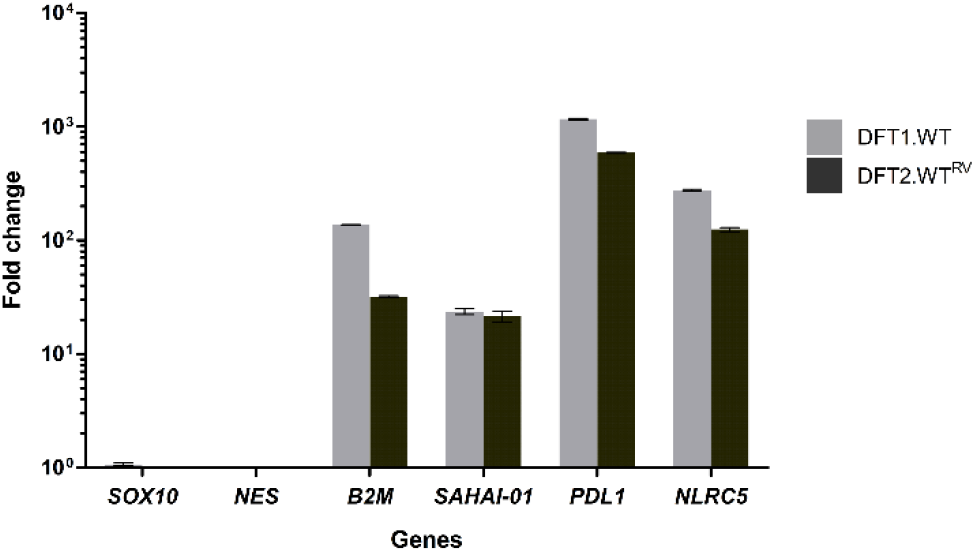
Upregulation of NLRC5 by IFNG in DFT1 and DFT2 cells. Fold change in mRNA expression (transcripts per kilobase million (TPM)) of *B2M*, MHC class I gene *SAHAI-01, PDL1* and *NLRC5* upon IFNG treatment in DFT1 C5065 cell line (DFT1.WT) and DFT2 RV cell line (DFT2.WT^RV^). *SOX10* and *NES* were included as internal controls. Bars show the mean of *N*=2 replicates per treatment. Error bars indicate standard deviation.

### NLRC5 upregulates MHC-I and antigen presentation genes but not PDL1 and non-classical MHC-I

To assess the role of NLRC5 in antigen processing and presentation, we developed an expression vector that stably upregulates NLRC5 in DFT cells. DFT1 cell line C5065 and DFT2 cell line JV were used for production of NLRC5-overexpressing DFT cells. Following drug selection to create stable cell lines, we performed RNA-seq on DFT1 and DFT2 cells stably transfected with BFP-control and NLRC5 vectors (see **Table 1** for list of cell lines). Changes in the mRNA expression profile of DFT cells overexpressing NLRC5 relative to BFP-control cells were examined in parallel with changes observed in wild-type DFT cells following IFNG treatment (**Fig. 2** and **Supplementary Fig. 3**). The transcriptome for IFNG-treated DFT2 cells was previously generated from the DFT2 RV cell line (DFT2.WT^RV^)^6^. Otherwise, all DFT2 results are from DFT2 JV.

**Figure 2.**
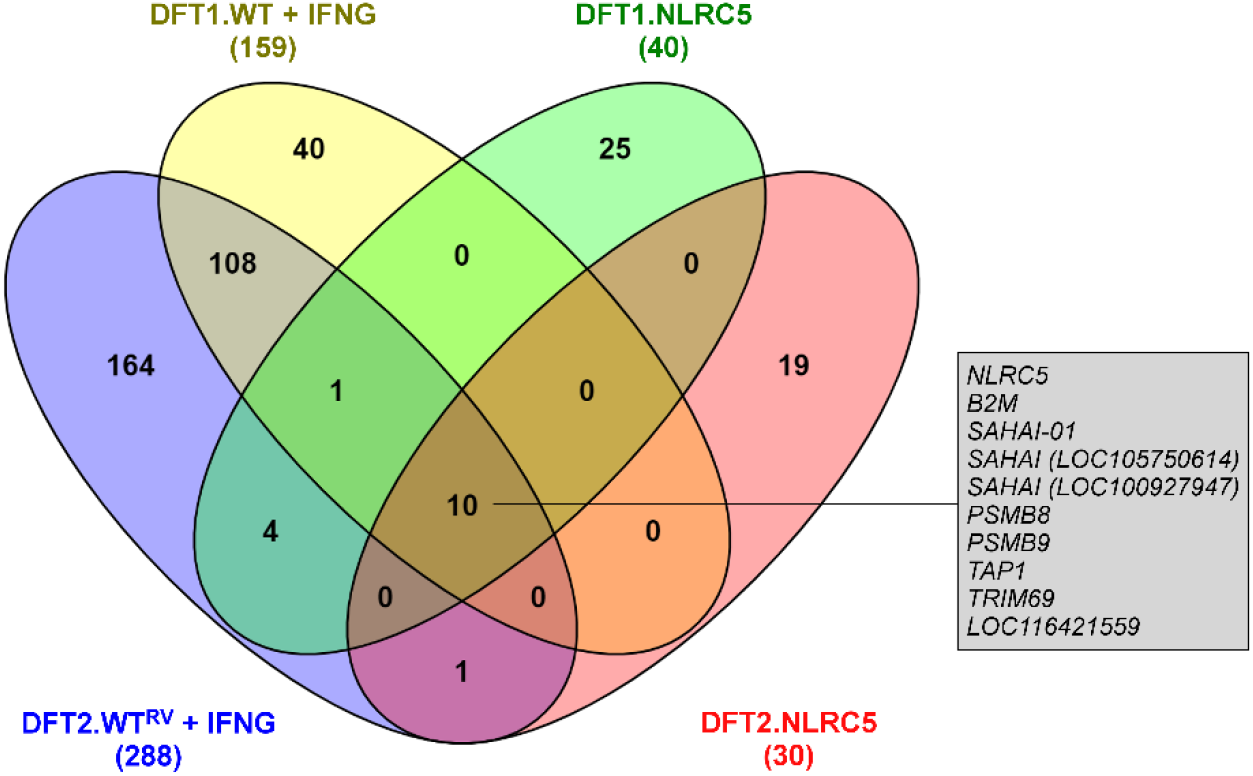
Venn diagram of genes significantly upregulated upon IFNG treatment and NLRC5 overexpression in DFT1 and DFT2 cells. Genes were defined as significantly upregulated when false discovery rate (FDR) < 0.05 and log_2_FC ≥ 2.0. Total number of genes upregulated for each treatment is indicated in parentheses under the sample name. The box shows genes upregulated in all four treatments: (i) IFNG-treated DFT1 cells (DFT1.WT + IFNG), (ii) IFNG-treated DFT2 cells (DFT2.WT^RV^ + IFNG), (iii) NLRC5-overexpressing DFT1 cells (DFT1.NLRC5), and (iv) NLRC5-overexpressing DFT2 cells (DFT2.NLRC5). See Supplementary Table 1 for a full list of differentially expressed genes and Supplementary Table 6 for description of devil-specific genes (LOC symbols).

Differential expression analysis showed that 159 genes were upregulated by IFNG (DFT1.WT + IFNG) in contrast to 40 genes by NLRC5 (DFT1.NLRC5) in DFT1 cells (**Fig. 2**). In DFT2 cells, 288 genes were upregulated by IFNG (DFT2.WT^RV^ + IFNG) and 30 genes by NLRC5 (DFT2.NLRC5) (**Fig. 2**). There were ten genes that were upregulated by both IFNG and NLRC5 in DFT1 and DFT2 cells. These shared genes were predominantly related to MHC-I antigen processing and presentation pathway which suggests a role of NLRC5 in IFNG-induced MHC-I expression.

A heatmap was used to explore the expression profiles of genes associated with MHC-I and MHC-II antigen processing and presentation. SOX10 and NES, which are Schwann cell differentiation markers highly expressed in DFT1 and DFT2 cells^6^, and the myelin protein periaxin (PRX), a marker for DFT1 cells^50^, were included as internal controls. Overall, NLRC5 upregulated genes involved in MHC-I antigen presentation to a smaller magnitude than IFNG (**Fig. 3**). NLRC5 upregulated a subset of IFNG-induced MHC-I genes *SAHAI-01*, *SAHAI (LOC105750614)* and *SAHAI (LOC100927947)*, and genes of the antigen processing machinery including *B2M*, *PSMB8*, *PSMB9*, and *TAP1*. In comparison, other IFNG-induced genes such as *PSMB10, TAP2* and TAP binding protein (*TAPBP*) were not upregulated by NLRC5 in either DFT1.NLRC5 or DFT2.NLRC5 cells. MHC-I genes that were induced by IFNG but not NLRC5 include non-classical MHC-I genes *SAHA-UK* and *SAHA-MR1*, although the latter was only induced in DFT2 cells treated with IFNG. Additionally, *PDL1* was upregulated by IFNG, but not NLRC5. Examination of the promoter elements immediately upstream of *SAHA-UK* and *PDL1* did not identify the putative MHC-I-conserved SXY module^51^ necessary for NLRC5-mediated transcription in the devil genome. A putative interferon-stimulated response element (ISRE) for devil MHC-I genes was identified 127 bp upstream of the start codon of *SAHA-UK* (**Supplementary Fig. 4**).

**Figure 3.**
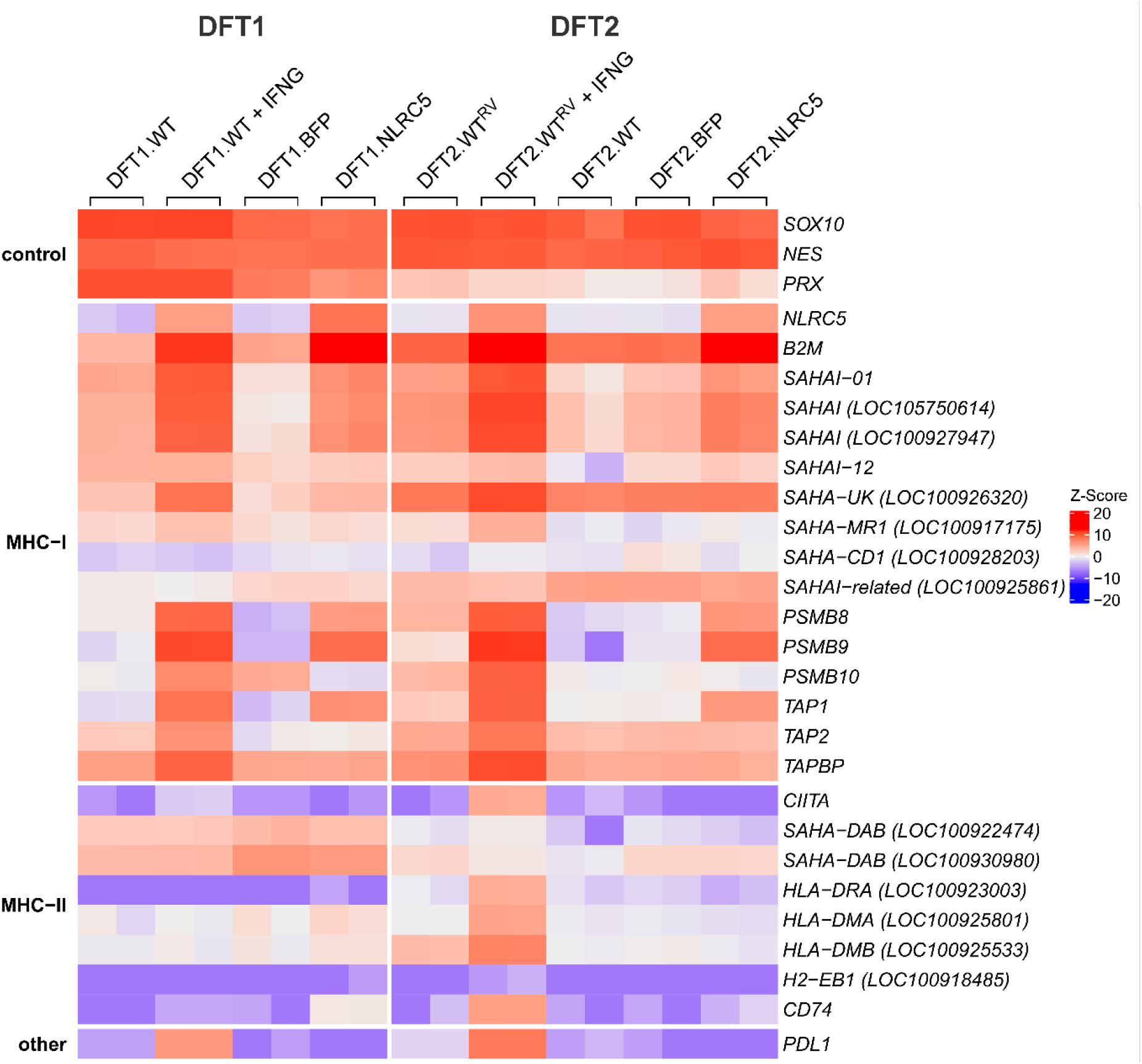
Heatmap showing expression profiles of genes involved in MHC-I and MHC-II antigen processing and presentation pathways, and PDL1 in IFNG-treated, and NLRC5-overexpressing DFT1 and DFT2 cells. Log_2_TPM expression values were scaled across each gene (rows) and represented by Z-Score, with red and blue representing high and low relative expression, respectively. Replicates for each treatment (*N*=2) are included in the heatmap. *SAHAI* encodes the Tasmanian devil MHC-I heavy chain gene. For genes with no official gene symbol (LOC symbols), alternative gene symbols were used according to the gene description on NCBI. See Supplementary Table 6 for corresponding NCBI gene symbols and description.

NLRC5 did not consistently regulate MHC-II genes. However, the invariant chain associated with assembly of MHC-II complexes, *CD74*, was significantly upregulated in DFT1.NLRC5. Similarly, IFNG treatment on DFT1 cells only upregulated MHC-II transactivator *CIITA*. Strikingly, IFNG treatment on DFT2 cells induced several MHC-II genes such as *HLA-DRA (LOC100923003), HLA-DMA (LOC100925801), HLA-DMB (LOC100925533), CD74* and *CIITA*.

### NLRC5 primarily functions in MHC-I antigen processing and presentation but is not limited to immune-related functions

The majority of research into NLRC5 has been devoted to its role as a regulator of MHC-I expression. In addition, some studies have reported possible roles of NLRC5 in antiviral immunity, inflammation and cancer through modulation of various signaling pathways^52-57^. To identify additional biological functions of NLRC5 in DFT cells, over-representation analysis of gene ontology (GO) biological processes and Reactome pathways was performed using the list of differentially expressed genes between NLRC5-overexpressing DFT cells and BFP-controls (FDR < 0.05, log_2_FC ≥ 2.0 or ≤ −2.0). Both analyses revealed significant up- and downregulation of genes associated with immune system processes and developmental processes in cells overexpressing NLRC5.

Among the list of genes upregulated in DFT1.NLRC5 and DFT2.NLRC5 cells, the most significantly associated GO biological process was *antigen processing and presentation of exogenous peptide antigen via MHC class I, TAP-dependent* (**Figs. 4A** and **5A**). Several additional immune-related processes were also associated with NLRC5 overexpression, particularly in DFT1 cells. Some of these included *positive regulation of immune response, interferon-gamma-mediated signaling pathway*, *immune response-regulating cell surface receptor signaling pathway* (**Fig. 4A**), and *regulation of interleukin-6 biosynthetic process* (**Fig. 4C**). In DFT1.NLRC5 and DFT2.NLRC5, GO terms related to development that were significantly over-represented included *morphogenesis of an epithelium* (**Fig. 4A**) and *negative regulation of epidermis development* (**Fig. 5A**), respectively.

**Figure 4.**
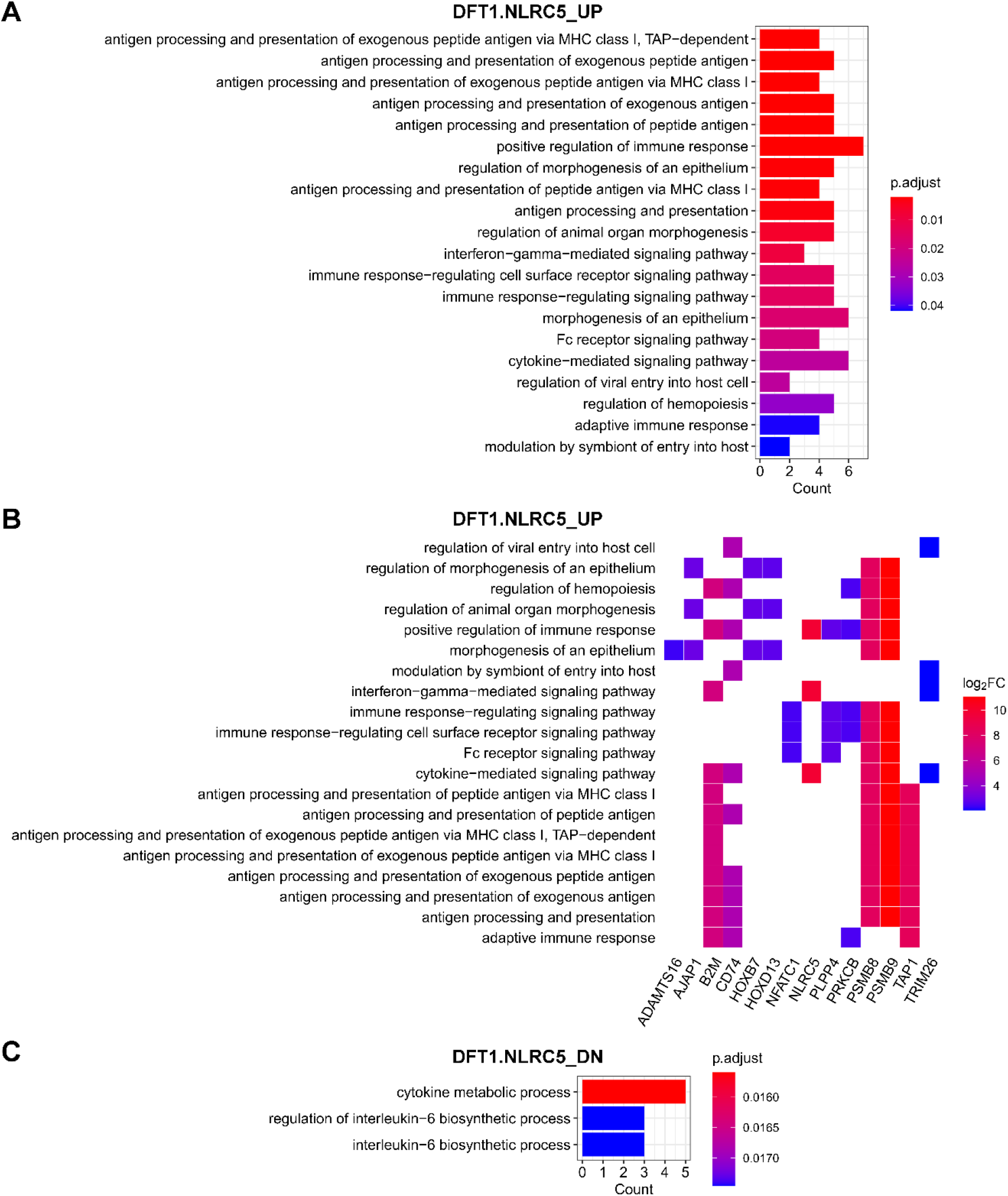
GO biological processes that were enriched in DFT1 cells with NLRC5 overexpression. GO biological process terms associated with genes upregulated (UP) **(A, B)** and downregulated (DN) **(C)** in DFT1.NLRC5. **(B)** Heatplot of genes associated with each positively-regulated GO term. The cut-offs p-value < 0.001 and adjusted p-value (p.adjust) < 0.05 were used to determine significant biological processes. P values were adjusted for multiple testing using Benjamini-Hochberg method. See also Supplementary Table 7 for full list of GO biological processes.

**Figure 5.**
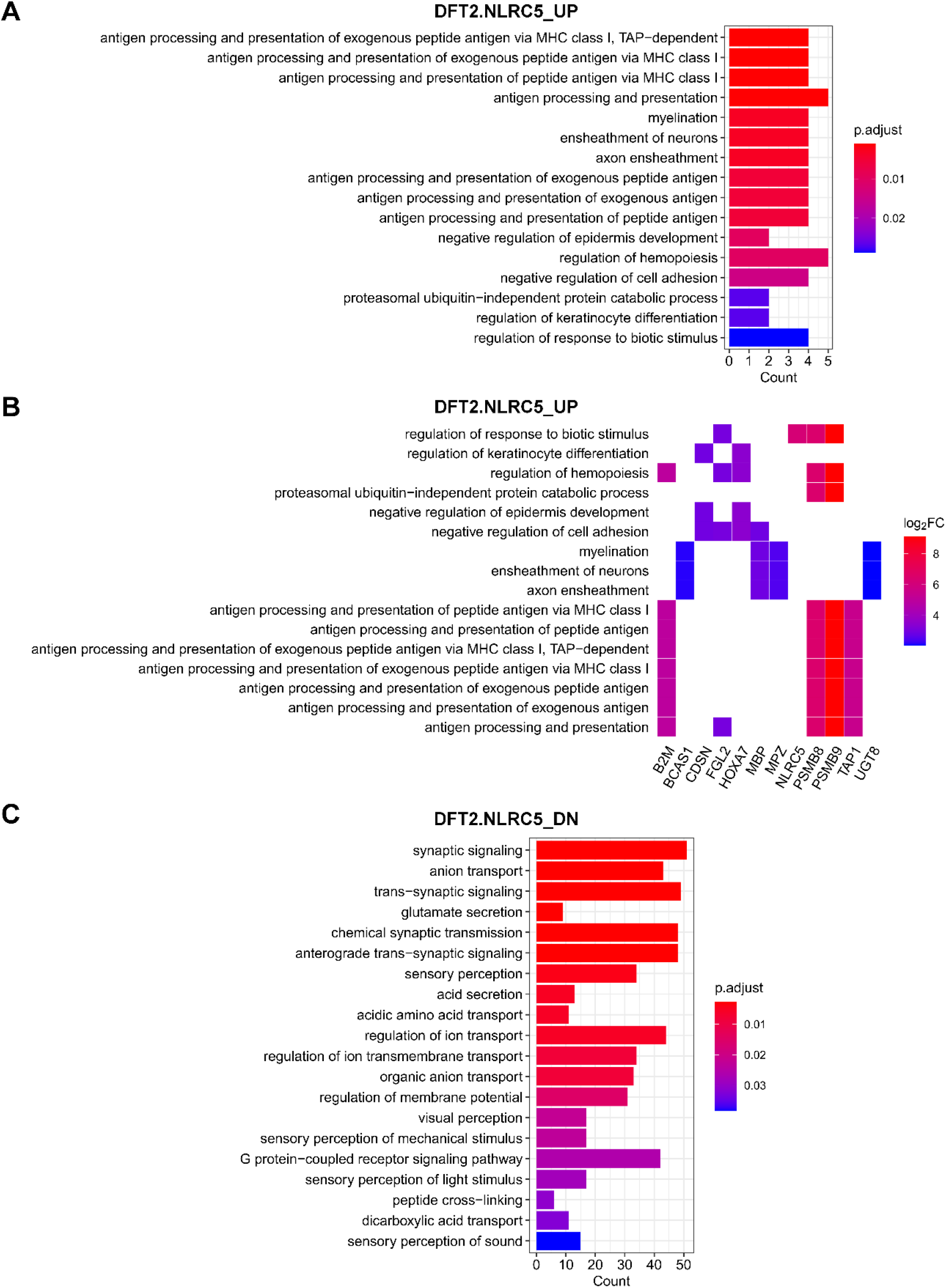
GO biological processes that were enriched in DFT2 cells with NLRC5 overexpression. GO biological process terms associated with genes upregulated (UP) **(A, B)** and downregulated (DN) **(C)** in DFT2.NLRC5. **(B)** Heatplot of genes associated with each positively-regulated GO term. The cut-offs p-value < 0.001 and adjusted p-value (p.adjust) < 0.05 were used to determine significant biological processes. P values were adjusted for multiple testing using Benjamini-Hochberg method. See also Supplementary Table 8 for full list of GO biological processes.

As DFT cells are of neuroendocrine origin, specifically of the Schwann cell lineage^5,6^, a number of neural-related genes were targeted by NLRC5. In DFT2 cells, NLRC5 upregulated genes that are involved in *myelination*, which are usually expressed at low levels in DFT2 cells^6^ (**Fig. 5A**). These genes include brain enriched myelin associated protein 1 (*BCAS1*), myelin binding protein (*MBP*), myelin protein zero (*MPZ*) and UDP glycosyltransferase 8 (*UGT8*) (**Fig. 5B**). Furthermore, many of the downregulated genes in DFT2.NLRC5 were related to nervous system function, mainly pertaining to *synaptic signaling* and *sensory perception* (**Fig. 5C**).

Reactome pathway analysis revealed an enrichment of pathways that were consistent with those identified by GO analysis. This included enrichment of the *ER-phagosome pathway* and *antigen processing-cross presentation* in DFT1.NLRC5 (**Table 2**) and DFT2.NLRC5 (**Table 3**); *signaling by the B cell receptor (BCR)* in DFT1.NLRC5; and *transmission across chemical synapses* in DFT2.NLRC5 cells. Interestingly, nuclear factor of activated T cells 1 (*NFATC1*), protein kinase C beta (*PRKCB), PSMB8* and *PSMB9*, associated with several GO immune-related processes in DFT1.NLRC5 (**Fig. 4B**), were enriched for the *beta-catenin independent WNT signaling* pathway (**Table 2**). Other enriched pathways included those involved in extracellular matrix organization such as *collagen chain trimerization* (**Table 2**) and *assembly of collagen fibrils and other multimeric structures* (**Table 3**).

**Table 2.** Reactome pathways enriched in differentially expressed genes in DFT1.NLRC5

| Reactome ID | Pathway | Count | Term size | p-value | p.adjust | Genes |
| --- | --- | --- | --- | --- | --- | --- |
| **Upregulated** |  |  |  |  |  |  |
| R-HSA-1236974 | ER-Phagosome pathway | 4 | 74 | 4.69E-05 | 4.75E-03 | *B2M, PSMB8, PSMB9, TAP1* |
| R-HSA-1168372 | Downstream signaling events of B Cell Receptor (BCR) | 4 | 80 | 6.37E-05 | 4.75E-03 | *NFATC1, PRKCB, PSMB8, PSMB9* |
| R-HSA-1236975 | Antigen processing-Cross presentation | 4 | 81 | 6.69E-05 | 4.75E-03 | *B2M, PSMB8, PSMB9, TAP1* |
| R-HSA-983705 | Signaling by the B Cell Receptor (BCR) | 4 | 104 | 1.77E-04 | 9.44E-03 | *NFATC1, PRKCB, PSMB8, PSMB9* |
| R-HSA-3858494 | Beta-catenin independent WNT signaling | 4 | 129 | 4.06E-04 | 1.73E-02 | *NFATC1, PRKCB, PSMB8, PSMB9* |
| R-HSA-1169091 | Activation of NF-kappaB in B cells | 3 | 64 | 7.19E-04 | 2.55E-02 | *PRKCB, PSMB8, PSMB9* |
| **Downregulated** |  |  |  |  |  |  |
| R-HSA-216083 | Integrin cell surface interactions | 5 | 62 | 1.16E-05 | 4.04E-03 | *CDH1, COL18A1, COL6A1, COL6A2, JAM2* |
| R-HSA-1251985 | Nuclear signaling by ERBB4 | 3 | 28 | 3.33E-04 | 4.93E-02 | *EREG, GFAP, S100B* |
| R-HSA-5173105 | O-linked glycosylation | 4 | 73 | 4.27E-04 | 4.93E-02 | *ADAMTS7, B3GNT7, GALNT13, GALNT17* |
| R-HSA-913709 | O-linked glycosylation of mucins | 3 | 34 | 5.96E-04 | 4.93E-02 | *B3GNT7, GALNT13, GALNT17* |
| R-HSA-8948216 | Collagen chain trimerization | 3 | 36 | 7.06E-04 | 4.93E-02 | *COL18A1, COL6A1, COL6A2* |

**Table 3.** Reactome pathways enriched in differentially expressed genes in DFT2.NLRC5

| Reactome ID | Pathway | Count | Term size | p-value | p.adjust | Genes |
| --- | --- | --- | --- | --- | --- | --- |
| **Upregulated** |  |  |  |  |  |  |
| R-HSA-1236974 | ER-Phagosome pathway | 4 | 74 | 3.43E-06 | 3.80E-04 | *B2M, PSMB8, PSMB9, TAP1* |
| R-HSA-1236975 | Antigen processing-Cross presentation | 4 | 81 | 4.93E-06 | 3.80E-04 | *B2M, PSMB8, PSMB9, TAP1* |
| R-HSA-983169 | Class I MHC mediated antigen processing & presentation | 5 | 312 | 5.96E-05 | 3.06E-03 | *B2M, PSMB8, PSMB9, TAP1, TRIM69* |
| R-HSA-983170 | Antigen Presentation: Folding, assembly and peptide loading of class I MHC | 2 | 18 | 3.31E-04 | 1.27E-02 | *B2M, TAP1* |
| R-HSA-162909 | Host Interactions of HIV factors | 3 | 119 | 6.91E-04 | 1.36E-02 | *B2M, PSMB8, PSMB9* |
| **Downregulated** |  |  |  |  |  |  |
| R-HSA-112316 | Neuronal System | 33 | 276 | 1.73E-06 | 1.28E-03 | *see Supplementary Table 10* |
| R-HSA-1474228 | Degradation of the extracellular matrix | 16 | 97 | 1.82E-05 | 6.73E-03 | *see Supplementary Table 10* |
| R-HSA-264642 | Acetylcholine Neurotransmitter Release Cycle | 5 | 10 | 5.87E-05 | 1.45E-02 | *see Supplementary Table 10* |
| R-HSA-181429 | Serotonin Neurotransmitter Release Cycle | 5 | 12 | 1.70E-04 | 2.27E-02 | *see Supplementary Table 10* |
| R-HSA-181430 | Norepinephrine Neurotransmitter Release Cycle | 5 | 12 | 1.70E-04 | 2.27E-02 | *see Supplementary Table 10* |
| R-HSA-112315 | Transmission across Chemical Synapses | 21 | 179 | 1.84E-04 | 2.27E-02 | *see Supplementary Table 10* |
| R-HSA-1474244 | Extracellular matrix organization | 24 | 224 | 2.65E-04 | 2.80E-02 | *see Supplementary Table 10* |
| R-HSA-166658 | Complement cascade | 6 | 21 | 4.02E-04 | 3.38E-02 | *see Supplementary Table 10* |
| R-HSA-1296072 | Voltage gated Potassium channels | 7 | 29 | 4.11E-04 | 3.38E-02 | *see Supplementary Table 10* |
| R-HSA-2022090 | Assembly of collagen fibrils and other multimeric structures | 9 | 49 | 5.62E-04 | 4.16E-02 | *see Supplementary Table 10* |
| R-HSA-210500 | Glutamate Neurotransmitter Release Cycle | 5 | 16 | 7.96E-04 | 4.91E-02 | *see Supplementary Table 10* |
| R-HSA-212676 | Dopamine Neurotransmitter Release Cycle | 5 | 16 | 7.96E-04 | 4.91E-02 | *see Supplementary Table 10* |

### NLRC5 induces MHC-I expression on the cell surface

To determine if NLRC5 is capable of regulating MHC-I expression at the protein level, surface MHC-I was analyzed by flow cytometry in DFT cells overexpressing NLRC5 using a monoclonal antibody against B2M^8^. The overexpression of NLRC5 induced upregulation of surface expression of B2M in both DFT1.NLRC5 (**Fig. 6A**) and DFT2.NLRC5 cells (**Fig. 6B**). The level of B2M expression was also comparable to wild-type DFT cells treated with IFNG.

**Figure 6.**
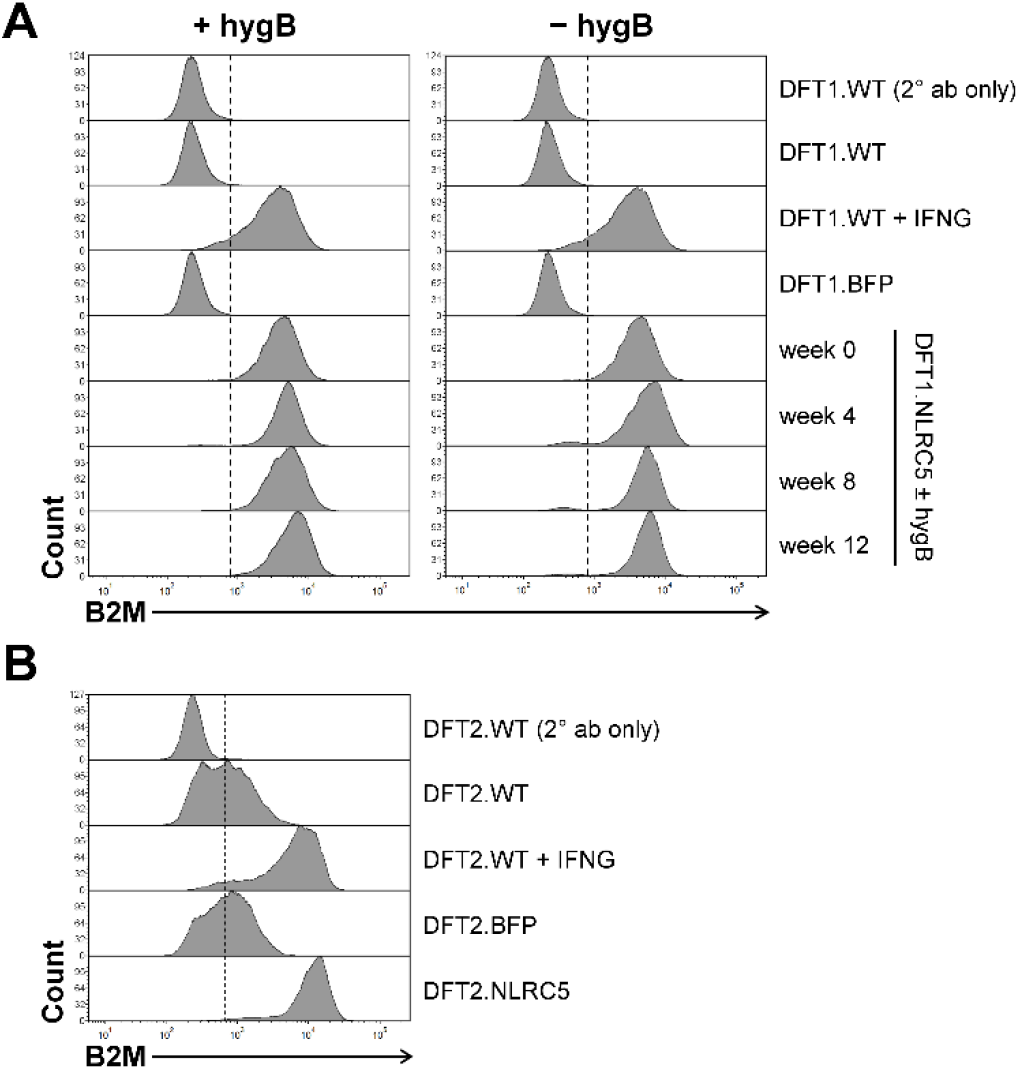
Upregulation of MHC-I following NLRC5 overexpression. Surface expression of B2M in DFT1.NLRC5 **(A)** and DFT2.NLRC5 **(B)**. B2M expression in the NLRC5 cell lines were compared to wild-type (DFT.WT), BFP-control (DFT.BFP), and IFNG-treated (DFT.WT + IFNG) DFT cells. **(A)** Stable expression of B2M in DFT1.NLRC5 was assessed every four weeks for 12 weeks post-drug selection in the presence and absence of hygromycin B (hygB) selection pressure. Secondary antibody-only staining (DFT.WT (2° ab only)) was included as a control. The results shown are representative of *N* = 3 replicates/treatment.

Next, we assessed the stability of NLRC5-induced MHC-I expression by examining the expression of B2M in long-term cultures. One-month post-drug selection, DFT1.NLRC5 cells cultured in the presence or absence of hygromycin B were stained for B2M every four weeks for a total of 12 weeks. As shown in **Fig. 6A**, MHC-I expression was stably maintained in DFT1.NLRC5 cells, with or without ongoing drug selection pressure throughout the 12-week culture thus, demonstrating the relative stability of the human EF1a promoter driving NLRC5 expression in long-term cell cultures. PDL1 was also not upregulated on the cell surface in NLRC5-overexpressing DFT cells compared to IFNG-treated DFT cells (**Supplementary Fig. 5**).

### MHC-I is a predominant target of anti-DFT antibody responses

It was previously reported that the antibodies from devils infected with DFT1 were specific to MHC-I, as determined by incubating serum from these devils with IFNG-treated DFT cells^12^. Considering the diverse roles of IFNG, there could be other IFNG-induced antigens that can serve as targets for the anti-DFT antibody response.

To establish if MHC-I is the target of anti-DFT serum antibodies, surface MHC-I expression was first ablated by knocking out the hemizygous *B2M* allele^10^ in wild-type DFT1 cells (DFT1.WT) and NLRC5-overexpressing DFT1 cells (DFT1.NLRC5) using CRISPR/Cas9 technology. Gene disruption of *B2M* was confirmed by genomic DNA sequencing (**Supplementary Fig. 2**), and flow cytometry using a monoclonal anti-B2M antibody (**Fig. 7**). CRISPR/Cas9-mediated *B2M* knockout (B2M^-/-^) in DFT1 cells rendered the cells irreversibly deficient for surface expression of B2M despite IFNG and NLRC5 stimulation (DFT1.B2M^-/-^ + IFNG and DFT1.NLRC5.B2M^-/-^). Due to the pivotal role of B2M in stability of MHC-I complex formation and surface presentation^58-62^, absence of surface B2M is indicative of a lack of surface MHC-I expression.

**Figure 7.**
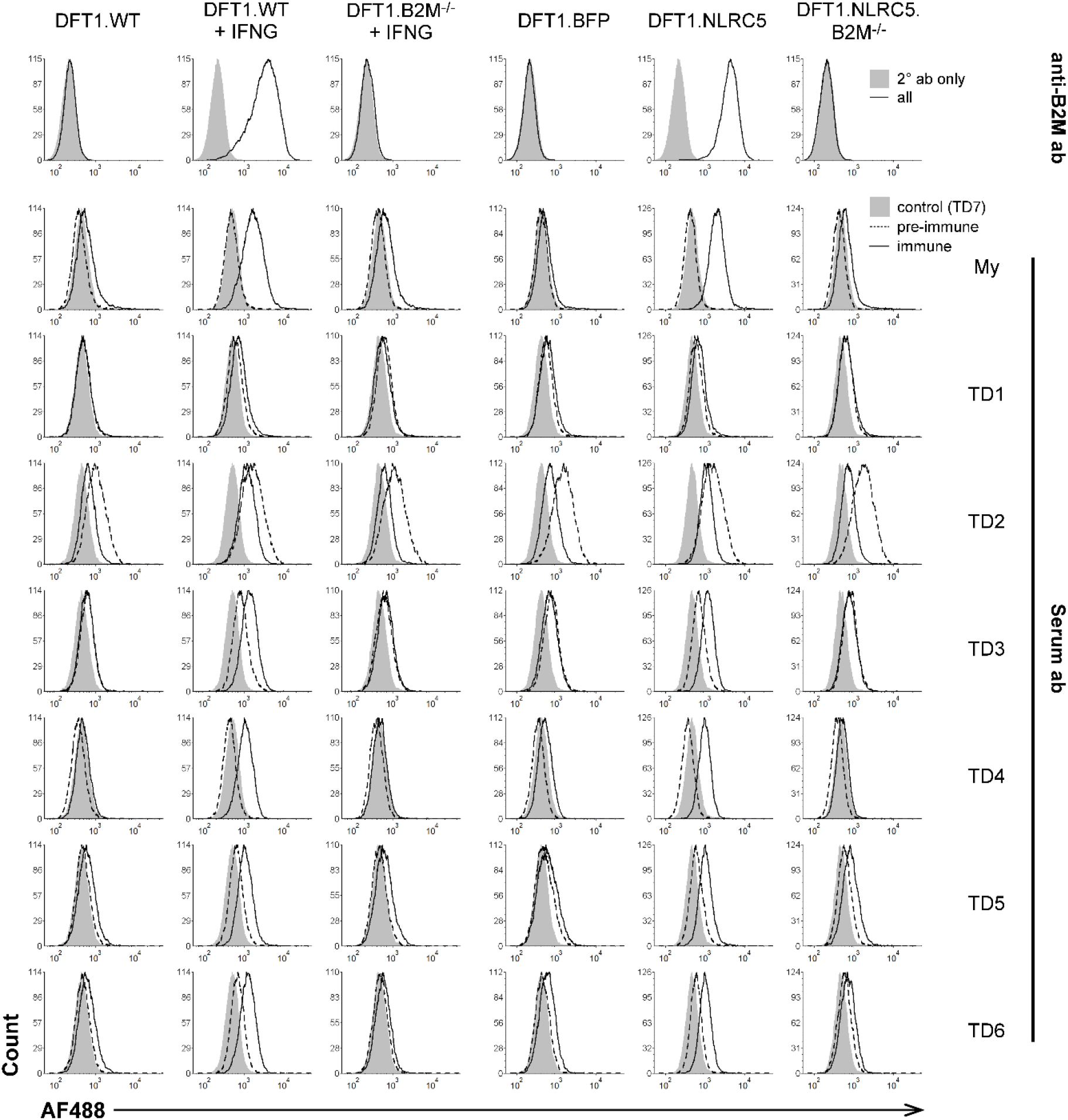
Flow cytometric analysis of serum antibody binding from devils with anti-DFT1 antibody response. Ablation of surface B2M in CRISPR/Cas9-mediated *B2M* knockout cells (B2M^-/-^) was confirmed using a monoclonal anti-B2M antibody (anti-B2M ab). Sera from six devils (TD1-TD6) with seroconversion (immune) following DFTD infection were tested against wild-type DFT1 (DFT1.WT), IFNG-treated DFT1 (DFT1.WT + IFNG), IFNG-treated *B2M* knockout DFT1 (DFT1.B2M^-/-^ + IFNG), BFP-control (DFT1.BFP), DFT1 overexpressing NLRC5 (DFT1.NLRC5) and *B2M* knockout NLRC5-overexpressing DFT1 (DFT1.NLRC5.B2M^-/-^) cells. An immunized devil with induced tumor regression (My) was included as a positive control, meanwhile serum from a healthy devil (TD7) was included as a negative control as represented in the shaded grey area. *Ab*, antibody; *AF488*, Alexa Fluor 488.

After surface MHC-I ablation was confirmed, serum from six wild devils (TD1–TD6) that demonstrated anti-DFT responses including natural DFT1 regressions^12^ was tested against *B2M* knockout cell lines DFT1.B2M^-/-^ and DFT1.NLRC5.B2M^-/-^. Serum from a healthy devil (TD7) and an immunized devil with induced tumor regression (My)^48^ were used as negative and positive controls for antibody binding. All six sera from DFTD^+^ devils (TD1-TD6) showed weak to no binding to DFT1.WT and DFT1.BFP, which are inherently negative for surface MHC-I (**Fig. 7**). With forced expression of MHC-I using IFNG (DFT1.WT + IFNG) and NLRC5 (DFT1.NLRC5), a positive shift in antibody binding was observed. There was no apparent difference in the level of antibody binding between IFNG-treated and NLRC5-overexpressing DFT1 cells, suggesting a similarity between the antibody target(s) induced by IFNG and NLRC5. Following *B2M* knockout, antibody binding of all six sera was reduced in both IFNG-induced (DFT1.B2M^-/-^ + IFNG) and NLRC5-induced *B2M* knockout DFT1 cells (DFT1.NLRC5.B2M^-/-^), suggesting that MHC-I is a target of DFT1-specific antibody responses in natural tumor regressions.

## DISCUSSION

Overexpression of NLRC5 in DFT cells has revealed a major and evolutionarily conserved role for NLRC5 in MHC-I antigen processing and presentation. Consistent with studies in human and mouse cell lines^9,18,63-65^, NLRC5 induced expression of classical MHC-I genes (*SAHAI-01, SAHAI* (*LOC105750614*), *SAHAI (LOC100927947)*), *B2M, PSMB8, PSMB9* and *TAP1* in both DFT1 and DFT2 cells. Despite the lack of increase in *TAP2* expression, the selective upregulation of MHC-I and other functionally-related genes by NLRC5 was sufficient to restore MHC-I molecules on the cell surface. Although the peptide transport function of TAP proteins typically involves the formation of TAP1 and TAP2 heterodimers, homodimerization of TAP proteins have been described^66,67^. However, the functionality of TAP1 homodimers remains to be verified. The conservation of genes of the MHC-I pathway, regulated by NLRC5 across species, highlights the important role of NLRC5 in MHC-I regulation.

Previous studies have shown that sera from wild devils with anti-DFT immune responses contained high titers of antibody that bound to IFNG-treated DFT1 cells. It was proposed that the primary antibody targets were MHC-I proteins^12^. Additionally, some of these devils experienced tumor regression despite the lack of strong evidence for immune cell infiltration into tumors. The function of NLRC5 that is mainly restricted to MHC-I regulation compared with IFNG provided an opportunity to re-examine antibody target(s) of serum antibodies from wild devils burdened with DFTs. A clear understanding of immunogenic targets of DFTs will provide direction for a more effective vaccine against DFTs.

The MHC-I complex was identified as the predominant target of anti-DFT serum antibodies. The antibody binding intensity against NLRC5-overexpressing DFT cells was similar to IFNG-treated DFT cells, suggesting similar levels of target antigen expression. When MHC-I expression was ablated through *B2M* knockout, antibody binding was reduced to almost background levels despite IFNG and NLRC5 stimulation. This discovery presents an option to exploit NLRC5 for induction of anti-DFT immunity via MHC-I expression, potentiated by the humoral anti-tumor response in Tasmanian devils. Although cellular immunity is likely a key mechanism for tumor rejection, B cells and antibodies can play eminent roles in transplant rejection^68^ and anti-tumor immunity^69^. B cells can promote rejection through antibody-dependent mechanisms that facilitate FcR-mediated phagocytosis by macrophages, antibody-dependent cellular cytotoxicity (ADCC) by NK cells, complement activation and antigen uptake by dendritic cells (reviewed by Yuen et al.)^70^. Moreover, B cells can enhance immune surveillance and response through direct antigen presentation to T cells and production of immune-modulating molecules such as cytokines and chemokines^70^.

Caldwell et al. reported that the most highly expressed MHC alleles on DFT2 cells are those that matched host MHC alleles^14^, which suggests that DFT cells may hide from host defenses or induce immunological tolerance via shared MHC alleles. If MHC-I is the major antibody target and potentially the overall immune system target, devils having the largest MHC mismatch with DFT cells will be the most likely to have strong MHC-I specific responses and reject DFTs, leading to natural selection in the wild. For example, previous studies have shown that some devils have no functional MHC-I allele at the UA loci and that these individuals can be homozygous at the UB and UC loci^48^. These individuals present a reduced MHC-peptide that would have the lowest probability of a match to the DFT MHC alleles that induce host DFT1 tolerance. However, selection for reduced genetic diversity in MHC alleles would be unfavorable for long-term conservation. A prophylactic vaccine would ideally be designed to assist in the preservation of the genetic diversity of wild devils^71^.

Although the MHC proteins themselves are likely a primary target of humoral and cellular immunity, MHC-I alleles generally differ by only a few amino acids^14,72^. Mutations in DFTs and somatic variation between host and tumor cells provide a rich source of additional antigenic targets for humoral and cellular immunity^10^. The reduction in antibody binding to *B2M* knockout cells suggests that these tumor antigens are unlikely to be the primary antibody targets, although binding of antibodies to peptide-MHC complexes cannot be excluded. Knocking out individual MHC alleles in DFT cells or overexpression of MHC alleles in alternative non-DFT cell lines could be used to disentangle the importance of specific alleles and investigate the potential for peptide-MHC complexes to be antibody targets.

Our results confirm that IFNG affects more immunoregulatory processes than NLRC5. However, the functional dichotomy of IFNG in cancer means that NLRC5 modulation could be an alternative to IFNG treatment for enhancing tumor cell immunogenicity in a range of species, including human. Importantly, NLRC5 upregulated B2M on the surface of DFT cells to similar levels as IFNG, but it does not upregulate inhibitory molecules. The restoration of functional MHC-I molecules without concomitant upregulation of PDL1 and SAHA-UK has multiple advantages over IFNG for triggering effective cytotoxic responses against DFT cells.

First, cells transfected with NLRC5 constitutively express MHC-I and therefore do not require culturing in IFNG, which can be problematic as IFNG can also reduce cell viability^17^. Second, PDL1 negatively regulates T cell responses by inducing T cell anergy^73^ and apoptosis^74^ while limiting T cell activity^75^. Moreover, PDL1 promotes tumor growth and survival by stimulating cell proliferation^76^ and resistance to T cell killing^77,78^. Third, the expression of monomorphic MHC-I SAHA-UK induced by IFNG would allow DFT cells to escape cytotoxic attack from both NK cells and CD8^+^ T cells^79^. Fourth, several other immune checkpoint protein receptor-ligand interactions were recently shown to be conserved in devils^80,81^, but we found no significant upregulation of these genes by NLRC5. The ability to improve tumor immunogenicity in the absence of inhibitory signals has positive implications for immunization and immunotherapeutic strategies. NLRC5 could evoke protective anti-tumor immunity against DFTs, similar to NLRC5-expressing B16-F10 melanoma cells in mice^64^.

The absence of a regulatory effect on *SAHA-UK* and *PDL1* by NLRC5 in contrast to IFNG could be due to the composition of the promoter elements of these genes. The promoter of MHC class I genes consists of three conserved cis-regulatory elements: a NFκB-binding Enhancer A region, an interferon-stimulated response element (ISRE) and a SXY module^82,83^. The SXY module is critical for NLRC5-mediated MHC-I transactivation as it serves as the binding site for the multi-protein complex formed between NLRC5 and various transcription factors^19,21,84^. An ISRE and SXY module is present within 200 base pairs of the start codon for all three classical devil MHC-I genes^51^. We identified an ISRE element in the *SAHA-UK* promoter region but were unable to identify an SXY module in this region. This could explain the upregulation of *SAHA-UK* upon IFNG stimulation but not in NLRC5-overexpressing DFT cells. Similarly, the SXY module was not identified in orthologues of *SAHA-UK*, which are *Modo-UK* in the grey short-tailed opossum^85^ and *Maeu-UK* in the tammar wallaby^86^. The difference in regulation and therefore, pattern of expression of the UK gene in marsupials^85-87^ may reflect a separate function from classical MHC-I. The marsupial UK gene has been hypothesized to play a marsupial-specific role in conferring immune protection to vulnerable newborn marsupials during their pouch life^86^. SXY modules are typically not found in the promoter region of PDL1^88^ therefore, it is not expected for NLRC5 to be a regulator of PDL1. Rather, IFNG-mediated induction of PDL1 occurs via transcription factor interferon regulatory factor 1 (IRF-1)^88^, which is induced by STAT1^89^.

Beyond MHC-I regulation, NLRC5 expression in DFT1 cells displayed other beneficial immune-regulating functions, mainly via the non-canonical β-catenin-independent WNT signaling pathway. One of the downstream effectors that was upregulated by NLRC5 included *PRKCB*, an activator of NFκB in B cells^90^. NFκB is a family of pleiotropic transcription factors known to regulate several immune and inflammatory responses including cellular processes such as cell proliferation and apoptosis^91^. In recent years, aberrations in NFκB signaling have been implicated in cancer development and progression^92,93^. The regulation of NFκB signaling by NLRC5 has been documented in several studies although the findings have been contradictory^54,55,94,95^.

In summary, we have demonstrated the role of NLRC5 in MHC-I regulation of DFT cells thereby, displaying the functional conservation of NLRC5 across species. The finding that MHC-I is a major antibody target in wild devils with anti-DFT response and natural DFT regression can help guide DFTD vaccine development and conservation management strategies. NLRC5-overexpressing DFT cells can be harnessed to elicit both cellular and humoral immunity against future and pre-existing DFT infections in wild devils using MHC-I as a target. Given the prevalence of altered MHC-I expression in cancer as a form of immune escape mechanism^96-98^, NLRC5 presents as a new target for providing an insight into the role of MHC-I in cancer as well as transplantation, and its manipulation for human cancer treatment and transplant tolerance.

## ACKNOWLEDGEMENTS

The authors would like to thank Patrick Lennard, Peter Murphy, and Candida Wong for assistance in the lab and Terry Pinfold for assistance in flow cytometry. We thank Hannah Siddle for supplying the monoclonal antibody for B2M and for offering her expertise in devil MHC-I immunogenetics. We wish to thank G. Ralph for ongoing care of Tasmanian devils, the Bonorong Wildlife Sanctuary for providing access to Tasmanian devils, and R. Pye for providing care for devils and collecting blood samples. This work was supported by ARC DECRA grant # DE180100484 and ARC Discovery grant # DP180100520, University of Tasmania Foundation Dr. Eric Guiler Tasmanian Devil Research Grant through funds raised by the Save the Tasmanian Devil Appeal (2013, 2015, 2017).

## AUTHOR CONTRIBUTIONS

ABL, ALP, ASF, CEBO and GMW designed the study. ALP, ASF, CEBO, GSL, JC, and JMD developed the technology. CEBO and JMD performed the experiments. ALP and CEBO performed bioinformatic analyses. CEBO created the figures. ALP, ASF, and CEBO analyzed the data. CEBO wrote the manuscript, and all authors edited the manuscript.

