## Supplementary Figure 1 for "NLRC5 regulates expression of MHC-I and provides a target for anti-tumor immunity in transmissible cancers" (from another version of this preprint: version 2, 2020-11-02)

**Andrew S. Flies, PhD**

Menzies Institute for Medical Research, College of Health and Medicine
University of Tasmania
Private Bag 23, Hobart TAS 7000

**Chrissie E. B. Ong, PhD candidate**

Menzies Institute for Medical Research, College of Health and Medicine
University of Tasmania
Private Bag 23, Hobart TAS 7000


**Supplementary Methods 1.** pAF105 plasmid construction.

**Supplementary Methods 2.** Flow cytometric analysis of PDL1 surface expression.

**Supplementary Table 1.** Differential expression analysis upon IFNG and NLRC5 induction in DFT cells [separate file].

**Supplementary Table 2.** PCR primers and reaction conditions.

**Supplementary Table 3.** Sequencing primers and reaction conditions.

**Supplementary Table 4.** Complementary oligonucleotides encoding the sgRNA sequence in *B2M*-targeting CRISPR/Cas9 vectors.

**Supplementary Table 5.** Serum sample information.

**Supplementary Table 6**. Alternative gene symbols used and the corresponding NCBI gene symbol.

**Supplementary Table 7.** Gene ontology (GO) biological processes that were enriched in DFT1 cells with NLRC5 overexpression [separate file].

**Supplementary Table 8.** Gene ontology (GO) biological processes that were enriched in DFT2 cells with NLRC5 overexpression [separate file].

**Supplementary Table 9.** Reactome pathways that were enriched in DFT1 cells with NLRC5 overexpression [separate file].

**Supplementary Table 10.** Reactome pathways that were enriched in DFT2 cells with NLRC5 overexpression [separate file].

**Supplementary Figure 1.** Schematic diagram of plasmids pAF105, pSBbi-BH and pCO1.

**Supplementary Figure 2.** Schematic diagram of CRISPR/Cas9 target sites and induced mutation in the devil *B2M* gene.

**Supplementary Figure 3.** Venn diagram of genes significantly downregulated upon IFNG treatment and NLRC5 overexpression in DFT1 and DFT2 cells.

**Supplementary Figure 4.** Promoter elements of devil MHC-I genes *SAHAI-01* and *SAHA-UK*.

**Supplementary Figure 5.** PDL1 and mTagBFP expression in DFT.NLRC5 cells.

**References**

**Supplementary Methods 1**

**pAF105 plasmid construction**

Mononuclear cells from devil lymph node were isolated and stimulated with 10 ng/mL recombinant devil IFNG[1] for 24 h to induce the expression of NLRC5. cDNA was extracted and the coding sequence for full length devil *NLRC5* (ENSSHAT00000015489.1) was isolated by PCR using primers deNLRC5.FOR and deNLRC5.REV (**Supplementary Table 2**). The isolated sequence was cloned into a tetracycline-controlled vector pTet-DualOFF (Clontech Laboratories, Mountain View, CA, USA) that was linearized by PCR (see **Supplementary Table 2** for primers and reaction conditions). NEBuilder® HiFi DNA Assembly Cloning Kit (NEB) was used to ligate the two fragments. The assembled plasmid pAF105 was transformed into NEB® 5-alpha competent *Escherichia coli* (High Efficiency) (NEB) according to manufacturer’s instructions. Positive clones were identified by colony PCR and the plasmid was purified using NucleoSpin® Plasmid EasyPure kit (Macherey-Nagel, Düren, Germany). The cloned devil *NLRC5* transcript was verified by Sanger sequencing using Big Dye™ Terminator v3.1 Cycle Sequencing Kit (Applied Biosystems (ABI), Foster City, CA, USA) and Agencourt® CleanSEQ® (Beckman Coulter, Brea, CA, USA) per manufacturer’s instructions. The sequences were analyzed on 3500xL Genetic Analyzer (ABI) (see **Supplementary Table 3** for list of sequencing primers).

**Supplementary Methods 2**

**Flow cytometric analysis of PDL1 surface expression**

Cells were harvested and plated in a round-bottom 96-well plate ($1\times10^5$ per well) and centrifuged at 500*g* for 3 min at 4 °C to discard the medium. Cells were blocked with 50 µL of 1% normal goat serum (Thermo Fisher Scientific) in FACS buffer (PBS with 0.5% BSA, 0.02% sodium azide) for 10 min on ice. After blocking, the cells were incubated with 50 µL of 2 µg/mL mouse anti-devil PD-L1 clone 1F8 antibody[1] labeled with DyLight 650 for 15 min on ice in the dark. The antibody was labeled using DyLight™ 650 Microscale Antibody Labeling Kit (Thermo Fisher Scientific). The cells were washed with 150 µL FACS buffer and centrifuged at 500*g* for 3 min at 4°C. Lastly, the cells were resuspended in 200 µL propidium iodide (PI) (500 ng/mL) (Sigma-Aldrich) prior to analysis on BD FACSCanto™ II (BD Biosciences, Franklin Lakes, NJ, USA). As a positive control for surface PDL1 expression, DFT1 C5065 and DFT2 JV cells were stimulated with 5 ng/mL recombinant devil IFNG[1] for 72 h.

**Supplementary Table 2.** PCR primers and reaction conditions.

| Vector | Template DNA | Target DNA | Product size (bp) | Primers | Primer sequence (5' to 3') | Reaction conditions |
|---|---|---|---|---|---|---|
| pAF105 | pTet-DualOFF | PCR-linearized pTet-DualOFF | 4267 | pAF105vec.FOR | CAGGATCCCAGACTGAACTTCTCCTAAGCGGCCGCATCGATAAGC | 1x: 95°C for 3 min;<br>35x: 98°C for 15 s, 68°C for 15 s, 72°C for 5 min;<br>1x: 72°C for 5 min |
| | | | | pAF105vec.REV | CTCATCATTCTGCCAGCTTAGGGAGTCCATGGTTGTGGCCATATTATCATCGTG | |
| | IFNG-stimulated devil PBMC cDNA | devil *NLRC5* cDNA | 5682 | deNLRC5.FOR | ATGGACTCCCTAAGCTGGCAGAATGATGAGGAAAGC | 1x: 95°C for 3 min;<br>35x: 98°C for 30 s, 68°C for 30 s, 72°C for 6 min;<br>1x: 72°C for 7 min |
| | | | | deNLRC5.REV | TTAGGAGAAGTTCAGTCTGGGATCCTGGCCCTGGAG | |
| pCO1 | pSBbi-BH | SfiI-digested pSBbi-BH | 6839 | N/A | N/A | N/A |
| | pAF105 | devil *NLRC5* cDNA | 5744 | pCO1.FOR | GTGAAAACTACCCCAAGCTGGCCTCTGAGGCCATGGACTCCCTAAGCTGGCAGAATG | 1x: 98°C for 30 s;<br>35x: 98°C for 10 s, 72°C for 30 s, 72°C for 3 min;<br>1x: 72°C for 2 min |
| | | | | pCO1.REV | AATTGATCCCCAAGCTTGGCCTGACAGGCCTTAGGAGAAGTTCAGTCTGGGATCCTGG | |

**Supplementary Table 3.** Sequencing primers and reaction conditions.

| Template DNA | Target DNA | Primers | Primer sequence (5' to 3') | Reaction conditions |
|---|---|---|---|---|
| pAF105 | devil *NLRC5* cDNA | deNLRC5_seq1.FOR | GCTGGAGTTGGTCAGAAAGT | 1x: 96°C for 1 min;<br>25x: 96°C for 10 s, 50°C for 5 s, 60°C for 4 min |
| | | deNLRC5_seq2.FOR | CCAGCTGCGGATTGAGAATTA | |
| | | deNLRC5_seq3.FOR | CTCAGACCTATCTGTCCTTCCT | |
| | | deNLRC5_seq4.FOR | GAGTCACCACCTTGGAGATTAG | |
| | | deNLRC5_seq5.FOR | TGCAGCCATCTTGACTTATCC | |
| | | deNLRC5_seq6.FOR | CTGGGTGATGAAGGACTCAAA | |
| | | deNLRC5_seq7.FOR | CTTAGCTGGTCCCAGGTTTAC | |
| | | deNLRC5_seq8.FOR | CAGGTGGAGGAGTTACACTTAG | |
| pCO1 | devil *NLRC5* cDNA | deNLRC5_seq1.FOR | GCTGGAGTTGGTCAGAAAGT | |
| | | deNLRC5_seq2.FOR | CCAGCTGCGGATTGAGAATTA | |
| | | deNLRC5_seq3.FOR | CTCAGACCTATCTGTCCTTCCT | |
| | | deNLRC5_seq4.FOR | GAGTCACCACCTTGGAGATTAG | |
| | | deNLRC5_seq5.FOR | TGCAGCCATCTTGACTTATCC | |
| | | deNLRC5_seq6.FOR | CTGGGTGATGAAGGACTCAAA | |
| | | deNLRC5_seq7.FOR | CTTAGCTGGTCCCAGGTTTAC | |
| | | deNLRC5_seq8.FOR | CAGGTGGAGGAGTTACACTTAG | |
| | | pSB_EF1a_seq.FOR | GCCTCAGACAGTGGTTCAAAG | |
| | | pSB_bGH_seq.REV | AGGCACAGTCGAGGCTGAT | |
| pAF217 | B2M_sgRNA_1 | U6 SeqFWD | GAGGGCCTATTTCCCATGATTC | 1x: 96°C for 1 min;<br>25x: 96°C for 10 s, 50°C for 5 s, 60°C for 75 s |
| pAF218 | B2M_sgRNA_2 | | | |
| DFT1.B2M$^{-/-}$ gDNA | *B2M* gene | B2m_seq1.FOR | AGTTTCCAAGCAAACGAAAATG | 1x: 96°C for 1 min;<br>25x: 96°C for 10 s, 50°C for 5 s, 60°C for 4 min |
| DFT1.NLRC5.B2M$^{-/-}$ gDNA | | B2m_seq1.REV | AGCTCTGGCTCCCAGAC | |

**Supplementary Table 4.** Complementary oligonucleotides encoding the sgRNA sequence in *B2M*-targeting CRISPR/Cas9 vectors.

| Vector | sgRNA | Target sequence (5' to 3') | Complementary oligonucleotides | Oligonucleotide sequence (5' to 3') |
|---|---|---|---|---|
| pAF217 | B2M_sgRNA1 | TCTTGCTAATTGCCCTCTTG | B2M_sgRNA_1.FWD | CACCGTCTTGCTAATTGCCCTCTTG |
| | | | B2M_sgRNA_1.REV | AAACCAAGAGGGCAATTAGCAAGAC |
| pAF218 | B2M_sgRNA2 | GATGCACAATTGCCCCAAGA | B2M_sgRNA_2.FWD | CACCGGATGCACAATTGCCCCAAGA |
| | | | B2M_sgRNA_2.REV | AAACTCTTGGGGCAATTGTGCATCC |

**Supplementary Table 5.** Serum sample information.

| Devil name | Devil ID | Sample collection date | |
|---|---|---|---|
| | | **Pre-immune** | **Immune** |
| TD1 | WPP75 | Nov 2009 | Nov 2011 |
| TD2 | WPP69 | May 2009 | Nov 2011 |
| TD3 | WPP287 | Feb 2011 | Feb 2014 |
| TD4 | WPP314 | July 2011 | May 2014 |
| TD5 | WPP209 | Nov 2010 | May 2011 |
| TD6 | WPP240 | Nov 2013 | Feb 2014 |
| TD7* | WPP213 | Feb 2012 | N/A |
| My[†] | Missy | 10 Mar 2011 | 3 Jan 2013 |

TD1–TD6 corresponds to TD1–TD6 in Pye et al.[2], My corresponds to TD1-My in Tovar et al.[3].

*TD7 was disease-free therefore, had no antibodies against either $\text{MHC-I}^-$ or $\text{MHC-I}^+$ DFT1 cells.

[†]My was an immunized devil with induced regression and had antibodies against $\text{MHC-I}^+$ DFT1 cells.

**Supplementary Table 6.** Alternative gene symbols used and the corresponding NCBI gene symbol.

| Alternative gene symbol | NCBI gene symbol | NCBI gene description | Also known as | Location in paper |
|---|---|---|---|---|
| PDL1 | CD274 | CD274 molecule | PDL1 | Figs. 1, 3 |
| LOC116421559 | LOC116421559 | uncharacterized | N/A | Fig. 2 |
| SAHAI (LOC105750614) | LOC105750614 | class I histocompatibility antigen, Gogo-OKO alpha chain | SAHA-I | Figs. 2, 3 |
| SAHAI (LOC100927947) | LOC100927947 | class I histocompatibility antigen, Gogo-OKO alpha chain | SAHA-I | Figs. 2, 3 |
| SAHA-UK (LOC100926320) | LOC100926320 | DLA class I histocompatibility antigen, A9/A9 alpha chain | N/A | Fig. 3 |
| SAHA-MR1 (LOC100917175) | LOC100917175 | major histocompatibility complex class I-related gene protein | MR1 | Fig. 3 |
| SAHA-CD1 (LOC100928203) | LOC100928203 | antigen-presenting glycoprotein CD1d | N/A | Fig. 3 |
| SAHAI-related (LOC100925861) | LOC100925861 | major histocompatibility complex class I-related gene protein | N/A | Fig. 3 |
| SAHA-DAB (LOC100922474) | LOC100922474 | DLA class II histocompatibility antigen, DR-1 beta chain | SAHA-DAB | Fig. 3 |
| SAHA-DAB (LOC100930980) | LOC100930980 | rano class II histocompatibility antigen, A beta chain | SAHA-DAB | Fig. 3 |
| HLA-DRA (LOC100923003) | LOC100923003 | HLA class II histocompatibility antigen, DR alpha chain | N/A | Fig. 3 |
| HLA-DMA (LOC100925801) | LOC100925801 | HLA class II histocompatibility antigen, DM alpha chain | N/A | Fig. 3 |
| HLA-DMB (LOC100925533) | LOC100925533 | HLA class II histocompatibility antigen, DM beta chain | N/A | Fig. 3 |
| H2-EB1 (LOC100918485) | LOC100918485 | H-2 class II histocompatibility antigen, E-D beta chain | N/A | Fig. 3 |

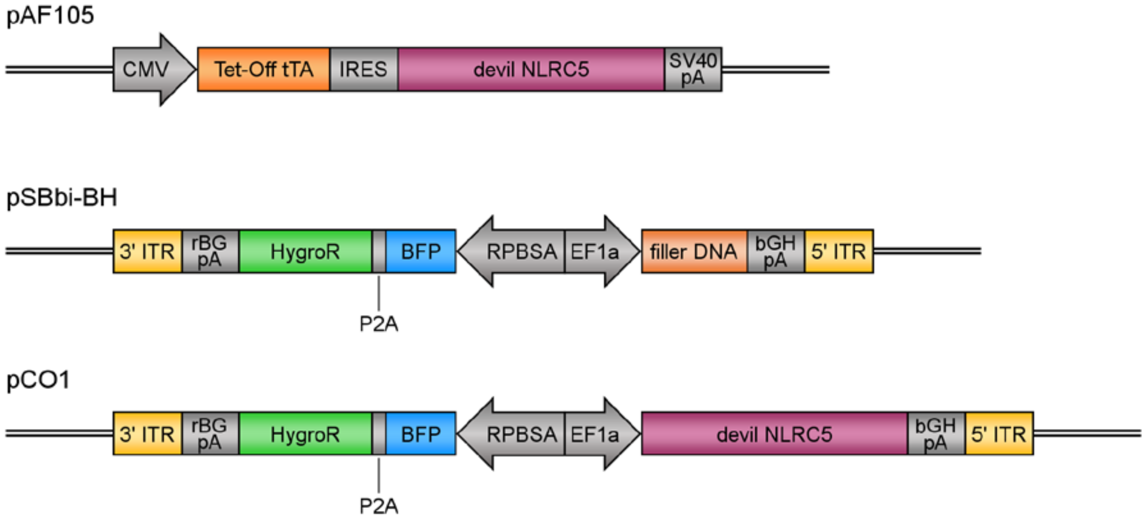


**Supplementary Figure 1. Schematic diagram of plasmids pAF105, pSBbi-BH and pCO1.** *Tet-Off tTA*, tetracycline(tet)-controlled transactivator; *IRES*, internal ribosomal entry site; *pA*, poly(A) tail; *ITR*, inverted terminal repeats; *HygroR*, hygromycin resistance gene; *BFP*, mTag-BFP.

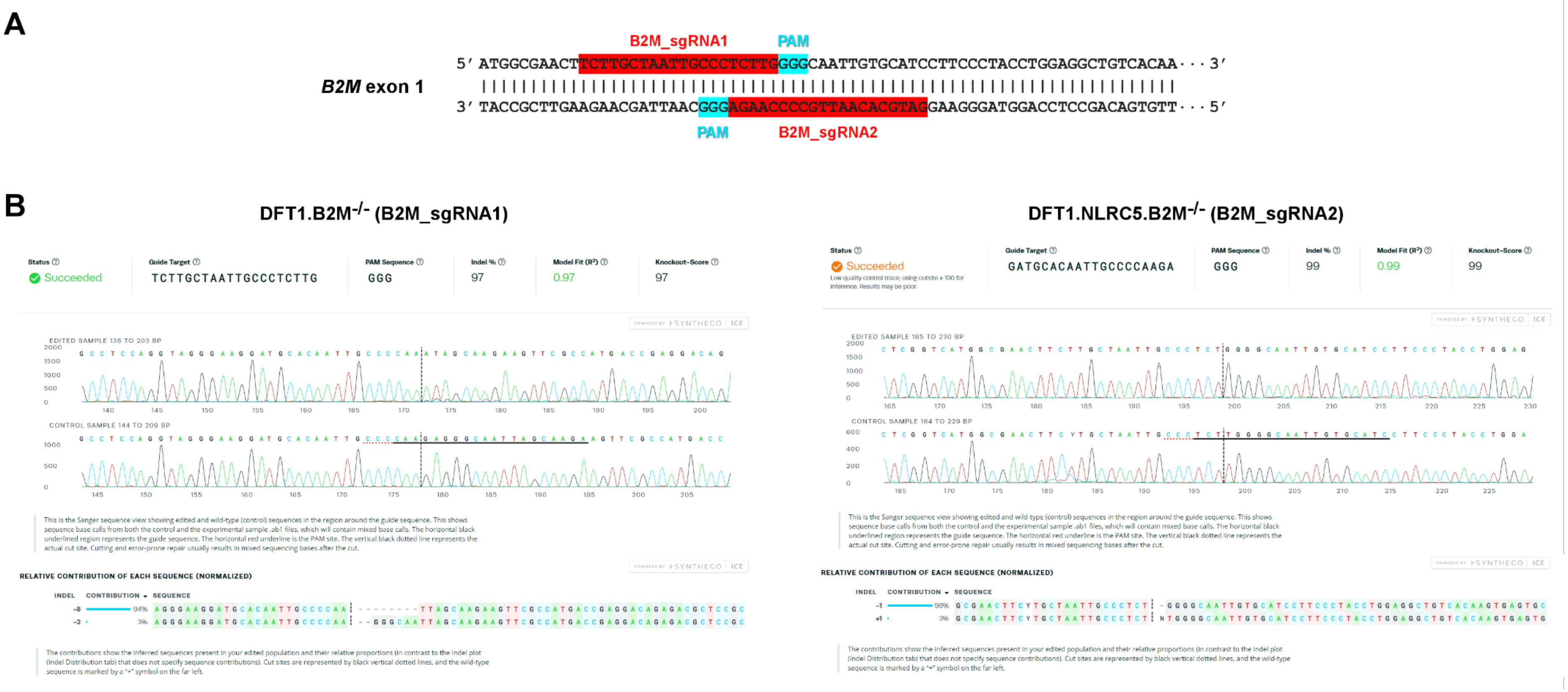


**Supplementary Figure 2. Schematic diagram of CRISPR/Cas9 target sites and induced mutation in the devil *B2M* gene. (A)** sgRNA target sequence and PAM sites on devil *B2M* exon 1. **(B)** Indel mutations and percentage, and knockout-score in *B2M* gene of FACS-sorted knockout cell lines DFT1.B2M$^{-/-}$ and DFT1.NLRC5.B2M$^{-/-}$. Indel percentage represents the percentage of sequences with edited traces (non-wild type sequence) compared to the control trace (wild-type sequence). Knockout score represents the proportion of cells that have frameshift mutations or more than 21 bp indels. 94% of DFT1.B2M$^{-/-}$ cells had an eight bp deletion and 3% had a two bp deletion in the *B2M* gene. In DFT1.NLRC5.B2M$^{-/-}$, 96% of the population had a one bp deletion and 3% had a one bp insertion in the *B2M* gene.

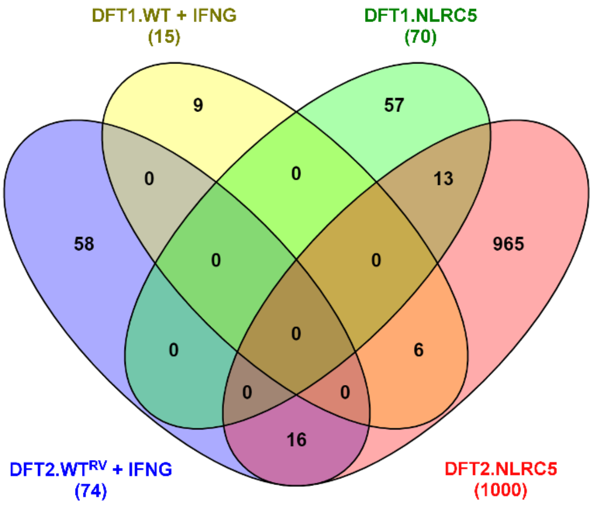


**Supplementary Figure 3. Venn diagram of genes significantly downregulated upon IFNG treatment and NLRC5 overexpression in DFT1 and DFT2 cells.** Genes were defined as significantly downregulated when false discovery rate (FDR) < 0.05 and $\log_2$FC $\leq -2.0$. Total number of genes downregulated for each treatment is indicated in parentheses under the sample name. See Supplementary Table 1 for a full list of differentially expressed genes.

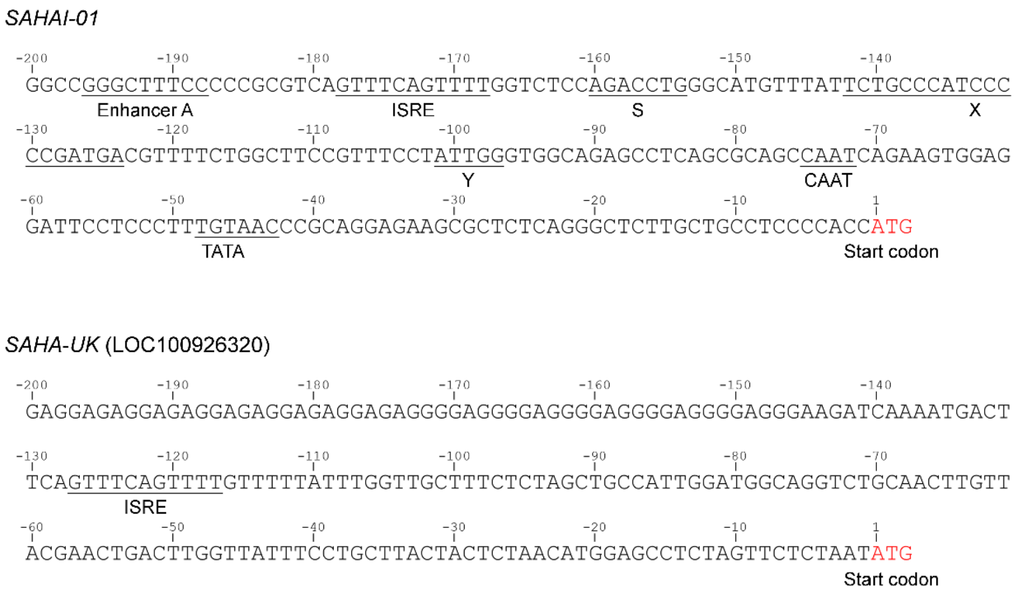


**Supplementary Figure 4. Promoter elements of devil MHC-I genes *SAHAI-01* and *SAHA-UK*.** Putative sites of cis-regulating promoter elements enhancer A, interferon-stimulated response element (ISRE), SXY module, CAAT and TATA boxes in devil classical MHC-I gene *SAHAI-01*[4] that were not identified within 1500 bp upstream of the start codon of non-classical MHC-I *SAHA-UK* with the exception of an ISRE element. The diagram above only shows the genomic sequence 200 bp upstream of the start codon.

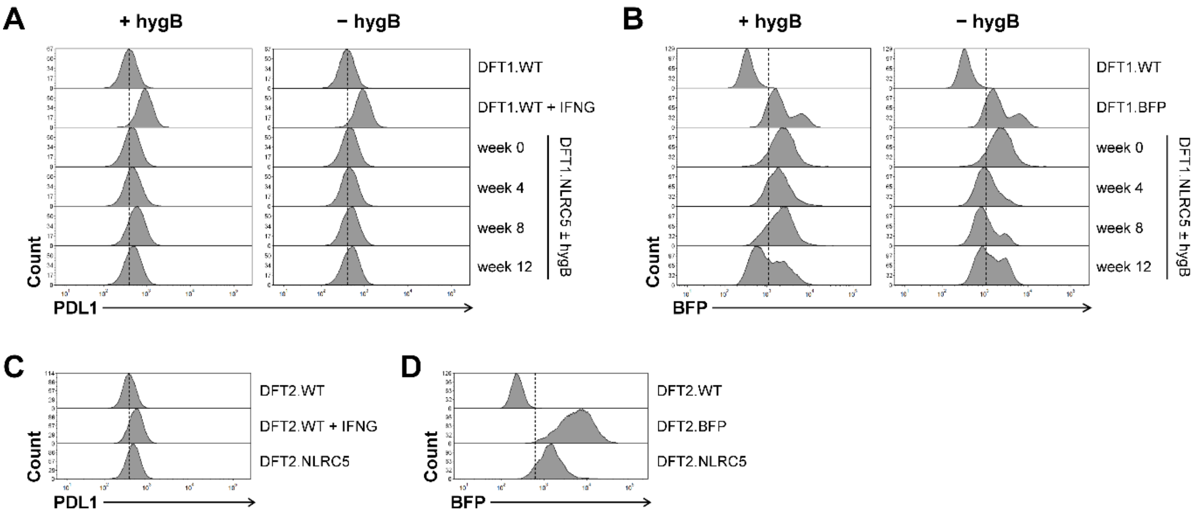


**Supplementary Figure 5. PDL1 and mTagBFP expression in DFT.NLRC5 cells. (A, B)** Expression of PDL1 and mTagBFP (BFP) in the presence and absence of hygromycin B (hygB) selection pressure in DFT1.NLRC5 cells, assessed every four weeks for 12 weeks post-drug selection. **(C, D)** Expression of PDL1 and mTagBFP in DFT2.NLRC5 cells. **(A, C)** PDL1 expression in the NLRC5 cell lines were compared against wild-type (DFT.WT) and IFNG-treated (DFT.WT + IFNG) DFT cells. *DFT.BFP*, DFT cells transfected with BFP-control vector. The results shown are representative of N = 3 replicates/treatment.
